# Evolving together: Cassandra retrotransposons gradually mirror promoter mutations of the 5S rRNA genes

**DOI:** 10.1101/2023.07.14.548913

**Authors:** Sophie Maiwald, Ludwig Mann, Sònia Garcia, Tony Heitkam

**Affiliations:** Faculty of Biology, Technische Universität Dresden, 01069 Dresden, Germany; Institut Botànic de Barcelona (IBB-CSIC), Passeig del Migdia s/n, 08038 Barcelona, Catalonia, Spain

**Author notes:** These authors share senior authorships and serve as communicating authors: Sònia Garcia, Tony Heitkam.

**Keywords:** 5S rDNA, ribosomal genes, 5S promoter, 35S-5S linkage, Cassandra, transposable elements, retrotransposons, long terminal repeats, concerted evolution, sequence mimicry, plant genomes, Asteraceae

## Abstract

The 5S rRNA genes are among the most conserved nucleotide sequences across all species. Similar to the 5S preservation we observe the occurrence of 5S-related non-autonomous retrotransposons, so-called Cassandra. Cassandras harbor highly conserved 5S rDNA-related sequences within their long terminal repeats (LTRs), advantageously providing them with the 5S internal promoter. However, the dynamics of Cassandra retrotransposon evolution in the context of 5S rRNA gene sequence information and structural arrangement are still unclear, especially: 1) do we observe repeated or gradual domestication of the highly conserved 5S promoter by Cassandras and 2) do changes in 5S organization such as in the linked 35S-5S rDNA arrangements impact Cassandra evolution? Here, we show evidence for gradual co-evolution of Cassandra sequences with their corresponding 5S rDNAs. To follow the impact of 5S rDNA variability on Cassandra TEs, we investigate the Asteraceae family where highly variable 5S rDNAs, including 5S promoter shifts and both linked and separated 35S-5S rDNA arrangements have been reported. Cassandras within the Asteraceae mirror 5S rDNA promoter mutations of their host genome, likely as an adaptation to the host’s specific 5S transcription factors and hence compensating for evolutionary changes in the 5S rDNA sequence. Changes in the 5S rDNA sequence and in Cassandras seem uncorrelated with linked/separated rDNA arrangements. We place all these observations into the context of angiosperm 5S rDNA-Cassandra evolution, discuss Cassandra’s origin hypotheses (single or multiple) and Cassandra’s possible impact on rDNA and plant genome organization, giving new insights into the interplay of ribosomal genes and transposable elements.

## Introduction

Despite both being repetitive genome components, the tandemly arranged ribosomal genes (rDNAs) and the dispersed transposable elements (TEs) seem to not have much in common. Yet, rDNAs and TEs can enter intricate co-evolution processes that are sometimes proposed, but neither fully appreciated nor understood; neither in their evolutionary mechanisms and implications nor in their effect on the genome.

Ribosomal RNAs (those encoded by ribosomal DNAs) comprise 80% of the RNA found in a typical cell and account for 60% of the ribosomal mass, having an essential role in protein synthesis (O’Connor and Adams 2010; Eaves et al. 2020). In eukaryotes there are four ribosomal RNA genes. The 18S, 5.8S, and 26S (28S/25S) rRNA genes are coded in a single operon (called 35S in plants and 45S rDNA in animals Hemleben et al. 2021) and the 5S rRNA gene is usually coded outside this operon, by 5S rDNA. These rDNAs are present in high numbers, from 50 to 13,000 copies per cell.

The 5S rDNA is perhaps the most enigmatic of the rRNA genes. It consists of a small transcriptional unit of about 120 base pairs and a non-transcribed spacer (NTS), normally clustered in long tandem arrays. While the 5S rRNA gene sequence is highly conserved, the NTS is variable in length and sequence, even in closely related species. The rapidly evolving nature of the NTS has been used for inferring interspecific relationships in many plant species (Perina et al. 2011; Waminal et al. 2014; Alexandrov et al. 2021). For such an apparently simple molecule of fundamental importance we know so very little, and many features of 5S remain unknown or controversial, e.g., although 5S rRNA is essential for the function of ribosomes, its specific role still remains unclear. Evolutionary biologists have conflicting views on its evolution, some assuming 5S rDNA to be a paradigm of concerted evolution, whilst others proposing alternative models (Brown et al. 1972; Nei and Rooney 2005). Another interesting feature is the range of genomic arrangements in which 5S rDNA can be found: although typically organized in tandems, it can be linked to their repetitive gene families (such as the 35S/45S DNA), scattered in the genome or located in linear or circular extrachromosomal DNA units (Drouin and de Sá 1995; Cohen et al. 2010; Rebordinos et al. 2013; Vierna et al. 2013). However, we still don’t know of any evolutionary advantage or biological functional differences of one 5S rDNA arrangement over another.

In contrast to the 5S rDNA, transposable elements (TEs) occur in a wide variety of structures and sequences (Bourque et al. 2018). Among those, long terminal repeat (LTR) retrotransposons are the most widespread in plants, sometimes accounting for more than 80 % of their genomes (Schnable et al. 2009). They are flanked by the name-giving LTRs that encode Polymerase II (Pol II) promoter motifs as well as transcription start and stop sites. Full-length retrotransposons usually carry all protein domains needed for their retrotransposition and can operate autonomously. Nevertheless, non-autonomous LTR retrotransposons also exist, with many not carrying any open reading frames at all.

Instead, these short terminal repeat retrotransposons in miniature (TRIMs) proliferate by exploiting the amplification machinery of their autonomous counterparts (Witte et al. 2001; Gao et al. 2016). Intriguingly, the ubiquitous TRIM family Cassandra deviates strongly from all other known TRIMs: Instead of carrying a retrotransposon-typical Pol II promoter, the Cassandra TRIMs have replaced this by co-opting the Polymerase III (Pol III) promoter from the 5S rRNA gene; hence, Cassandra retrotransposons and the 5S rDNA often share considerable sequence stretches and adopt similar secondary structures after transcription (Kalendar et al. 2008). Being unusually widespread across plant genomes (Gao et al. 2016), including monocots, dicots, and ferns (but not gymnosperms), Cassandras are believed to be ancient (Antonius-Klemola et al. 2006; Kalendar and Schulman 2006; Yin et al. 2014; Gao et al. 2016; Maiwald et al. 2021).

Both being ancient parts of the genome, actively using Pol III promoters and generating RNAs that adopt distinct secondary structures, Cassandra retrotransposons and 5S rDNAs share many similarities. Nevertheless, it is still unclear how Cassandra retrotransposons and the 5S rDNA depend on each other and if these two sequence classes co-evolve. We aim to understand how changes in the 5S rDNA sequence, structure and organization are mirrored by Cassandra retrotransposons.

To tackle this question, we focus on the Asteraceae, probably the largest and more diverse plant family, which harbors a wide variation in 5S rDNA sequence, structure and genomic organization. The Asteraceae family comprises over 25,000 species that fall into more than 1700 genera (Mandel et al. 2019). An unusual 5S rDNA organization, in which the 5S rDNA is linked to the 35S rDNA, has been detected in three large groups of subfamily Asteroideae (tribes Anthemideae, Gnaphalieae and the Heliantheae alliance), accounting for nearly 25% of this families’ species (Garcia et al. 2010). A later study detected a significant sequence divergence in the conserved C-box of the 5S promoter in some, but not all, Asteraceae with 35S-5S linkage (Garcia et al. 2012). Thus, the Asteraceae harbor a range of 5S rDNAs variations – much more than other plant families. If Cassandra retrotransposons are impacted by changes in the 5S rDNA, this plant family offers a superior starting point to understand any potential 5S rDNA-Cassandra co-evolution. To date, over 25 Asteraceae genome assemblies are available (www.plabipd.de; last accessed 22.05.2023), offering a generous resource for investigating the potential rDNA–retrotransposon co-evolution.

Here, we systematically follow Cassandra and 5S rDNA evolution across plants and especially within the Asteraceae. To better understand the make-up of canonical Cassandra retrotransposons and their dependence on the 5S rDNA, we first mined all published sequences across plants. Then, focusing on the Asteraceae with their diverging 5S rDNA landscapes, we analyzed 15 *de novo* identified Cassandra sequences from 22 Asteraceae genomes and checked how 5S rDNA changes may impact Cassandra evolution. For this, we targeted two major shifts in 5S rDNA evolution: Promoter sequence mutation and emergence of the 35S-5S linkage.

## Material and Methods

### Plant material and genomic DNA sequencing

Plants of *Artemisia annua* (MV8) from the living collection of the Institut Botànic de Barcelona (Spain), and *Tragopogon porrifolius* (TRA18) provided by the IPK Genebank of the Plant Genome Resources Center Gatersleben (Germany), were grown under long day conditions in the greenhouse. Genomic DNA was extracted from 1-3 g of fresh leaf material with a modified CTAB (cetyltrimethyl/ammonium bromide) protocol after (1987) and Cullings (1992).

For both species WGS library preparation (TruSeq DNA kit) and sequencing was carried out by Macrogen Inc. Europe, using an Illumina NovaSeq machine. The sequencing yielded around 4.5 Gb of 151 bp paired-end reads for each species, with an insert size of 660 bp (*A. annua*) and 470 bp (*T. porrifolius*), respectively.

### Retrieval of published plant Cassandra retrotransposon sequences

We manually extracted representative Cassandra retrotransposon sequences from all published reports that targeted plant genomes. For further analyses, we included 66 Cassandra-named sequences from these studies, which we could unambiguously determine to be Cassandra retrotransposons by manual annotation (supp. table 1).

### Screening of genome sequence assemblies for new Cassandra retrotransposons

As some Asteraceae species show an unusual variation in 5S rDNA sequence and genomic arrangement, we focused on *de novo* identification of Cassandra sequences in these species. For identification purposes we used published genomes of 22 Asteraceae species (supp. table 1). A multi-query BLAST search with known plant Cassandra sequences against these genomes was not successful. The first *de novo* identification in an Asteraceae genome was performed in *Artemisia annua* with TRIM-specific LTR Finder settings (Gao et al. 2016): -d 30, -D 2000, -l 30, -L 500 and relaxed parameters for motif detection, only looking for PBS and PPT motifs (no conserved regions, TG - CA architecture or TSD sequences to detect diversified sequences). LTR-Finder hits were extracted and used to perform a multi-query BLAST search (Megablast, wordsize 28, Gap cost: 2/2, scoring 1/-2) with known Cassandra sequences and the *Artemisia annua*-specific 5S rRNA gene. BLAST hits were processed by manual inspection, followed by annotation of a reference full length sequence. The *Artemisia annua*-specific Cassandra sequence was included in the Cassandra dataset, which was then again used for a multi-query BLAST against the remaining Asteraceae genome assemblies. Genomes of Cichorioideae subfamily showed no Cassandra positive BLAST hits and therefore were screened with the LTR-Finder routine again, but yielded no results.

### 5S rRNA gene retrieval and identification

5S rRNA gene information for selected species was obtained from the 5S rRNA database (http://combio.pl/rrna/Szymanski et al. 2016) and the NCBI nucleotide database. As we focused especially on species within the Asteraceae, of which some do not have published 5S rDNA data, we performed Readcluster analysis with the RepeatExplorer pipeline on Galaxy (Novák et al. 2013). Paired-end, randomly selected WGS datasets (Illumina) were downloaded from the ENA and analyzed by the pipeline RepeatExplorer (https://repeatexplorer-elixir.cerit-sc.cz/galaxy). For long read data we used the RepeatExplorer Utilitier “Create sample of long reads” and “Get pseudo short paired end reads from long reads” to adapt read length. After checking quality with FastQC (Andrews 2010), reads were trimmed and adapters removed when present. Read sampling (5 million reads per pair) was carried out, reads were interlaced and subsequently analyzed by TAREAN (a tool for the identification of genomic tandem repeats from NGS data), as implemented in RepeatExplorer (default settings). 5S rDNAs were detected as one of the tandem repeats present in the analyzed genomes and the consensus sequence was extracted for each of the analyzed species. These 5S rDNA sequences were compared with a standard 5S rDNA reference sequence (*Arabidopsis thaliana* E006, from the 5SrRNAdb by Szymanski et al. 2016) with Geneious Prime® 2023.0.1, and the corresponding genic portion of each of the target species was extracted.

### Determining the linkage between 5S rDNA and 35S rDNA by low coverage assembly

To confirm the potential linkage of the ribosomal genes WGS data from selected species (supp. table 1) was retrieved from the respective sequence archives (ENA, SRA, and GSA). The read quality was checked using FastQC 0.11.9 (Andrews 2010) and quality or adapter trimming was carried out using Trimmomatic 0.39 (Bolger et al. 2014) when necessary. Reads were sub-sampled to match a 1x genome coverage using seqtk 1.3 (Li et al. 2013) or RepeatExplorer Utilities “Create sample of long reads” (https://www.elixir-czech.cz/; Novák et al. 2020) for short and long reads, respectively. For sequencing data with less available data all available reads were used. For the first assembly round MEGAHIT 1.2.9 (Li et al. 2016) with the meta-large preset was used and only contigs larger than 5 kb were kept. The second assembly round was done with SPAdes genome assembler 3.15.5 (Bankevich et al. 2012) using the isolate preset, a coverage cut-off of 20 and the megahit final contigs as trusted contigs. The resulting assemblies were visualized using the Bandage 0.9.0 (Wick et al. 2015) assembly viewer. Nodes were colored by BLAST hits using the ribosomal genes of *Helianthus annuus* and drawn around the BLAST hits using a distance of 5-25, respectively.

### Cassandra metadata and comparison with 5S rRNA genes

Sequence comparison for Cassandra full length and LTR sequences was performed with multi sequence alignments (MUSCLE; Edgar 2004) and manual refinement. Cassandra sequences were then grouped into families according to their level of similarity. We applied a pairwise identity threshold of 70 % for family assignment. Exceptions were made for families with different variants. These families showed variants with indels, which affect the overall alignment pairwise identity in a negative way. But nevertheless, sequences could be grouped into one family due to their similarity in non-indel areas. To compare species-specific 5S rRNA genes with the corresponding Cassandra sequence we performed species-, lineage-, and family-specific alignments (MUSCLE) and dotplots.

All studies regarding nucleotide sequence motifs and variability of Cassandra sequences were performed by manual inspections of the MUSCLE alignments.

### Data availability

All accession numbers for published genome assemblies, Cassandra retrotransposons and 5S rRNA genes are listed in supp. table 1. *De novo* Cassandra sequences identified in this study are available at NCBI under the following study: PRJEB61458 (acc. numbers: OX591319-OX591330 and OY284499-OY284501). Sequence alignments for plant family specific Cassandra LTRs are available at Zenodo: https://zenodo.org/record/8144620. WGS raw sequence data of *Artemisa annua* and *Tragopogon porrifolius* data for this study have been deposited in the European Nucleotide Archive (ENA) at EMBL-EBI under accession number PRJEB63080 (https://www.ebi.ac.uk/ena/browser/view/PRJEB63080, ERR11535563 and ERR11535566).

## Results

### Cassandras form a lineage in plants: They share 5S promoters in their LTRs, but differ in sequence and length

To better understand the relationship between Cassandra retrotransposons and the 5S rDNA, we first compiled a Cassandra dataset across different plant species and expanded it with full length Cassandras from published Asteraceae genomes. Our motivation to investigate the structural hallmarks of Cassandra retrotransposons beyond the Asteraceae were initial difficulties in Cassandra identification: As Cassandras carry highly conserved regions and share LTR similarities across the plant kingdom (Maiwald et al. 2021), one could assume that they constitute a single Cassandra family with derivatives, scattered across plant genomes. However, for Asteraceae genomes, an initial similarity search via BLAST was not successful, indicating either greater divergence in Cassandra sequence or absence in Asteraceae genomes. Adopting a *de novo* approach relying solely on structure-based criteria, however, we retrieved Cassandra sequences for 15 out of the 22 investigated Asteraceae genome sequences. Sequence-wise, Asteraceae-derived Cassandras differ slightly from previously published Cassandra sequences (see below).

We complemented this Asteraceae dataset with 66 published Cassandra sequences from other plants, reaching 81 Cassandra reference sequences, each representing the Cassandra retrotransposon landscape of the respective genome. This dataset allows us to investigate structural hallmarks of Cassandra sequences across the plants and to understand how Asteraceae Cassandras compare to other plant Cassandras.

Being non-autonomous LTR retrotransposons, all Cassandras in our dataset show no remains of any coding regions. Sequence-wise, all of them show LTR/LTR identities of at least 74 % (suppl. table 2). Structurally, they harbor a methionine *primer binding site* (PBS) and a *polypurine tract* (PPT) for first and second strand synthesis by a reverse transcriptase (suppl. table 2).

Regarding element and LTR lengths, Cassandra sequences across the angiosperms show a wide range of sizes. The smallest Cassandra in our dataset is from *Saruma henryi* (Aristolochiaceae) with a length of 563 bp, whereas the largest from *Glycine max* (Fabaceae), with a length 968 bp, is almost double in size. It is noticeable that Cassandra full-length elements are more or less equal-sized within one plant family (supp. table 2). Furthermore, longer Cassandra full length retrotransposons also tend to have rather long LTR sequences. For example, the longest Cassandras reside within the Fabaceae (Cassandra length approx. 900 bp; LTR lengths: approx. 400 bp), whereas smaller representatives are found in the Piperales and Polypodiales (Cassandra length: approx. 600 bp; LTR length approx. 200 bp; figure 1, supp. table 2). Similar rules apply to the internal region. There seems to be a plant family-specific conserved preference of internal region size lengths, which, in contrast to the LTR lengths, does not correlate with the overall length.

**Figure 1:**
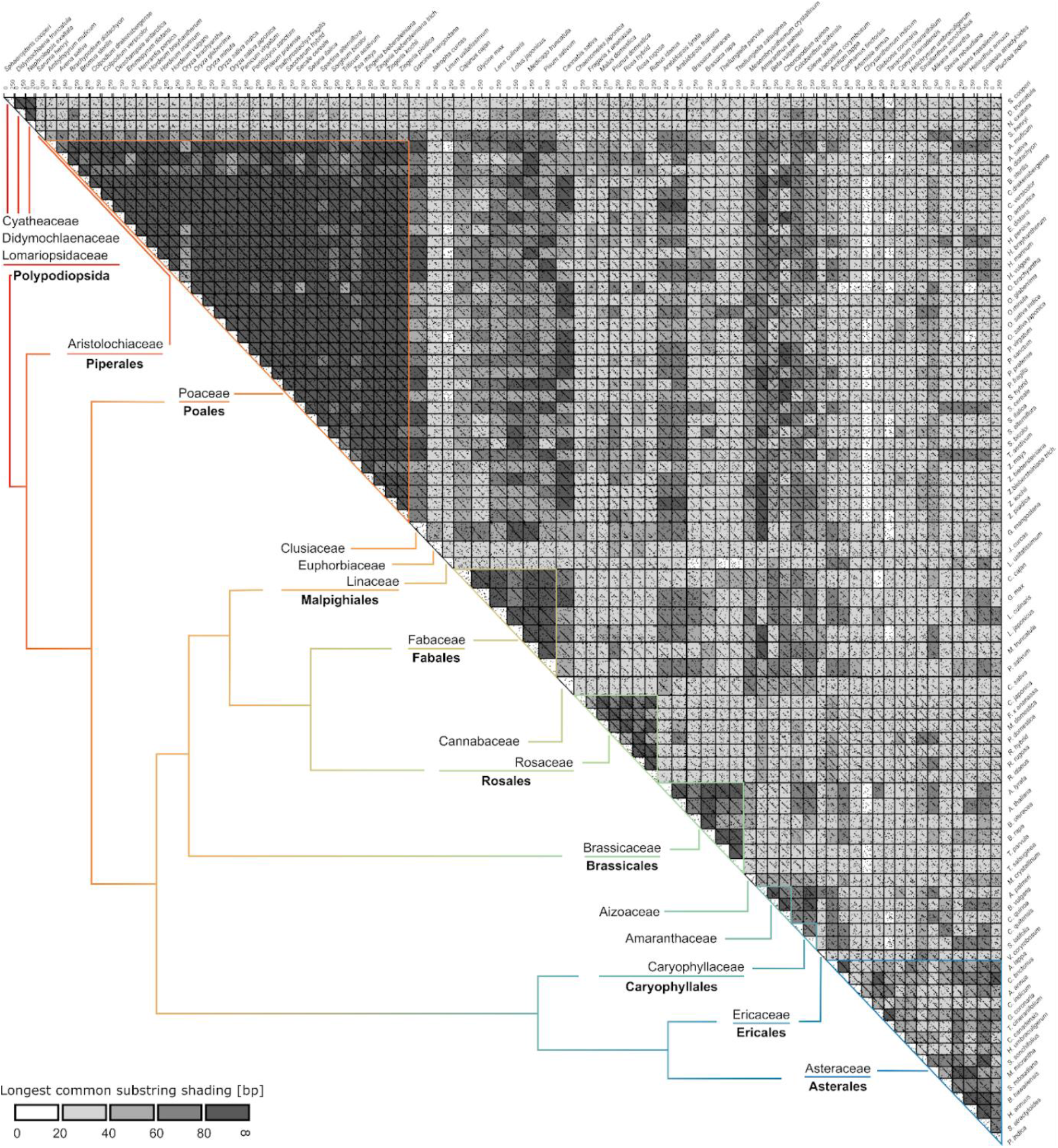
LTR comparison of 81 Cassandra retrotransposons. We performed an all-against-all dotplot analysis, where similarities between defined windows (k = 12, allowed mismatches n= 3) are marked as a dot. If sequences show larger regions of similarity, multiple dots form linear patterns. Sequences are shaded according to the longest common substring, meaning sequences with a high number of matching windows are shaded in dark gray and white/light gray for sequences with a small number of similar sequence units. Each and every sequence shows at least a small diagonal, which information-wise can be mainly limited to the conserved region Cassandra sequences share with the 5S rDNA and a short promoter-flanking region. Cassandra LTR sequences show increased similarity within one plant family but tend to accumulate mutations for more distal ones. Visualization of phylogenetic relationships are indicated by colored lines and resemble the current taxonomy proposed by the Angiosperm Phylogeny group (2017).

Placing the Asteraceae Cassandra retrotransposons into this framework of typical plant Cassandra lengths, they are on the smaller side, due to their short LTR and internal sequences with median length values of 270 bp and 82 bp respectively, similarly to Cassandras from the Rosaceae, Amaranthaceae and ferns.

Regarding LTR sequence conservation, a typical family-defining feature of an LTR retrotransposon, the variation is very high across all 81 plant Cassandra sequences. All of them have an overall pairwise identity of 44.4 %. Dotplot comparison about all plant Cassandras led us to the identification of Cassandra families based on plant family-specific similarities (see a comparative analysis among all Cassandra species in figure 1). A closer look into LTR identities also revealed high values > 70 % within different plant families (table 1; plots with darker shading in figure 1). The exceptionally low sequence identity for Cassandras within the Fabaceae and Asteraceae can be explained by variant formation (suppl. figure 1) due to indels within the LTRs of certain species.

To understand the main hallmarks of Cassandra retrotransposons and to assess the extent of Cassandra variability in angiosperms, we compared the sequence relationships of the Cassandra LTR regions, the internal regions, and the plant species. As seen in table 1, Cassandra shows high pairwise identity values >70 % for at least one of the two main sequence compartments: LTR (Poaceae, Brassicaceae, Caryophyllaceae, Amaranthaceae) or internal region (Fabaceae). Exceptions are Rosaceae Cassandras, with the highest values of pairwise identity for both LTR and internal region, and Asteraceae Cassandras, with the lowest conservation for both structural components. A patchy pattern for Asteraceae Cassandras is also seen within the dotplot (figure 1) and sequence alignments (suppl. figure 2) and resembles the phylogenetic assignment to the corresponding tribes of these species. Hence, a higher similarity of LTRs is not accompanied by conservation of the internal region and the other way around. Also, transcription-wise, there seems to be no need to preserve either LTR and/or internal regions as all combinations of similarity patterns in the two components are present across Cassandra families.

**Table 1:** Median length [bp] and family pairwise identity values [%] of plant Cassandras for families with two or more members. The number of representative Cassandras resembles the number of species with Cassandra retrotransposons within this plant family.

| plant family | # of representative Cassandras | median length [bp] | overall pairwise identity [%] | LTR pairwise identity [%] | internal pairwise identity [%] |
| --- | --- | --- | --- | --- | --- |
| Poaceae | 32 | 731 | 74.5 | 77.1 | 65.5 |
| Fabaceae | 6 | 912 | 56.0 | 55.3 | 75.1 |
| Rosaceae | 7 | 665 | 82.1 | 81.9 | 85.5 |
| Brassicaceae | 6 | 804 | 73.9 | 74.4 | 64.8 |
| Caryophyllaceae | 2 | 803 | 69.4 | 76.4 | 50.2 |
| Amaranthaceae | 3 | 761 | 64.2 | 74.7 | 42.4 |
| Asteraceae | 15 | 630 | 61.4 | 61.6 | 67.5 |

Considering this context, we find that LTRs of Asteraceae Cassandras are characterized by their short lengths (table1, suppl. table 2) and lesser conservation compared to most of the other plant families studied.

### Some Asteraceae harbor a Cassandra-related, non-autonomous LTR retrotransposon without the iconic 5S promoter

As we investigated BLAST results queried against the newly identified Asteraceae Cassandra from *A. annua*, we observed a mix of Cassandra-positive hits in the genomes of the Carduoideae. This led us to identify, for the first time, Cassandra-related non-autonomous LTR elements without the 5S promoter. These occur in some Asteraceae species of subfamily Carduoideae. These Cassandra-like elements share a part of their internal region with the canonical Cassandra, but differ in LTR sequence and length (suppl. figure 2). Most strikingly, as the iconic Cassandra 5S promoter is missing, these sequences do not represent Cassandra retrotransposons, but canonical terminal-repeat retrotransposons in miniature (TRIMs). In two species (*Arctium lappa* and *Carthamus tinctorius*) these Cassandra-like TRIMs co-exist with Cassandra, whereas in *Cynara cardunculus* only the Cassandra-like TRIM is present. As such, these Cassandra-like TEs may represent evolutionary progenitors or intermediates.

### Plant Cassandra sequences share a conserved 5S rDNA similarity region

Out of the 81 Cassandra retrotransposons in our dataset, we compared them to their corresponding published or newly annotated Asteraceae 5S rRNA gene, if available. By definition, all Cassandra sequences harbor a region similar to their corresponding 5S rRNA gene. The length of this similarity region is species-specific and always located approximately in the middle of the LTR sequence (suppl. table 2). The shared similarity region spans to the internal control region (ICR) with the promoter motifs (A-Box, intermediate element (IE) and C-Box) in all Cassandras. Although some species, like *Malus* × *domestica* (suppl. table 2), show a large similarity region of more than 100 nt, we were able to define a “core” unit of ∼ 70 bp present in all Cassandra sequences (suppl. table 2, figure 2). Although the core unit itself is present and conserved in all Cassandras, the position of the similarity regions within the LTR varies (suppl. table 2), but it is never located at the LTR termini. We observe family-specific similarity regions located within a region of 25 – 75 % of all LTR nucleotide positions.

**Figure 2:**
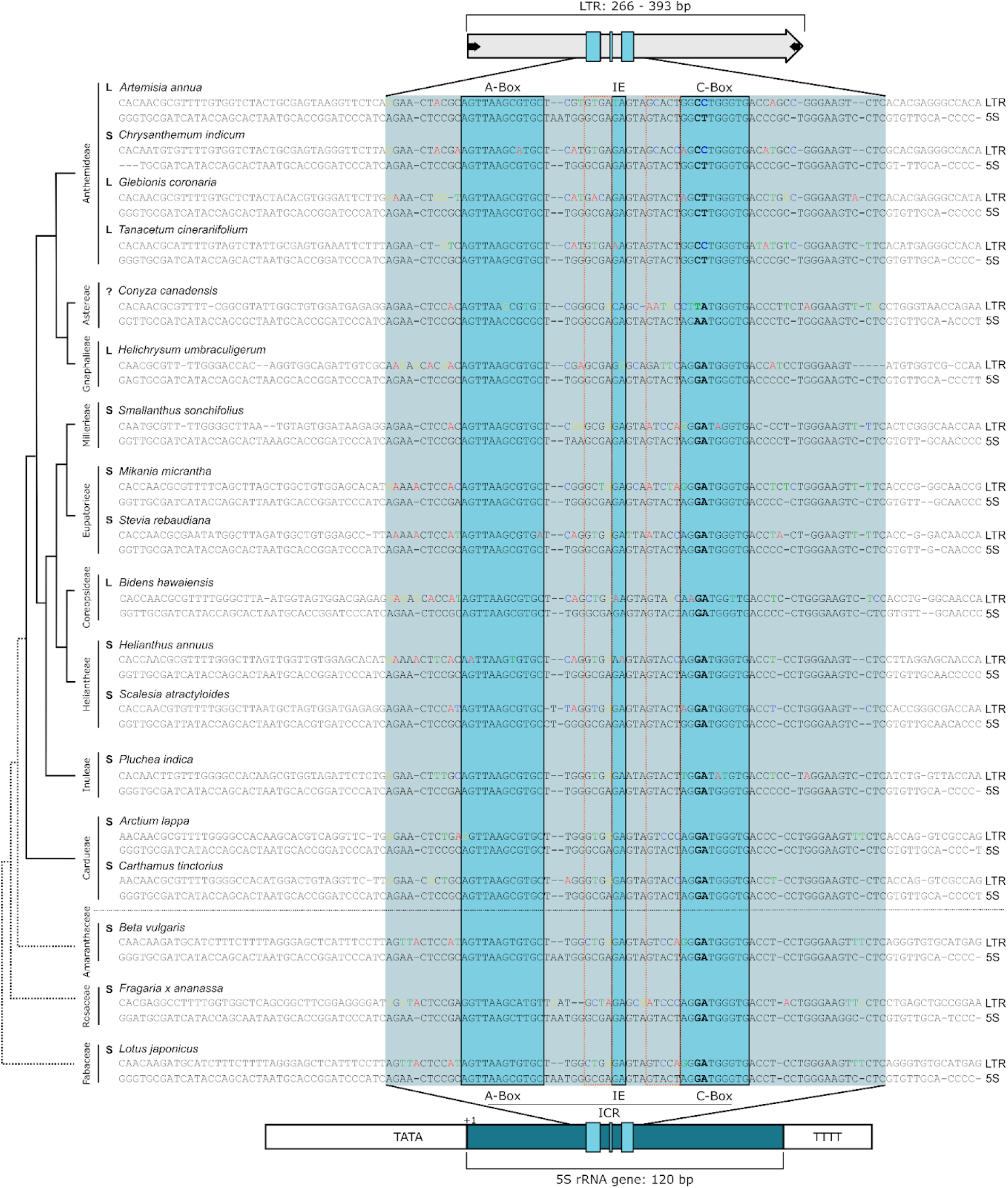
Similarity region shared by the 5S rRNA gene and Cassandra retrotransposons in the Asteraceae and other plants. Differences from the 5S gene are highlighted in colors (A = red, T = green, C = blue, G = yellow). Key sequence variation in the C-Box is highlighted in bold. The arrangement of the 5S rDNA in a linked (L) or separated (S) configuration is indicated (see also figure 3). The regions of high sequence divergence between the Box motifs are marked with an orange rectangle.

Within this highly conserved similarity region there is a very prominent variation: two kinds of C-Boxes. One of these motifs is solely specific for the Asteraceae Cassandra sequences within the Anthemideae tribe, the other is found in the remaining species. These prominent deviations can be explained by a closer look at the corresponding genomic 5S rRNA genes. In certain Asteraceae species, the usually highly conserved C-Box is slightly different, showing a 5’-GG**<u>CT</u>**TGGGTG-3’ motif instead of the canonical 5’-AG**<u>GA</u>**TGGGTG-3’. This C-Box variation is mirrored by the corresponding Cassandra sequence, although it is not identical, showing a 5’-GG**<u>CC</u>**TGGGTG-3’ motif (figure 2; bold bases within the C-Box). We hence observe that Cassandra elements somehow mimic changes in the 5S promoter of the 5S rRNA gene.

### The 5S promoter motif changed after the emergence of the 35S-5S linkage in the Asteraceae

Interestingly, the 5S promoters with the mutated C-Box emerged in the Anthemideae, a tribe well-known for harboring 5S genes in a linked arrangement, such as species of the genus *Artemisia* (Garcia et al. 2009). We wondered if there was any relation between the 35S-5S linkage and the promoter shift, and if they evolved independently or within similar timespans.

We already have data regarding the promoter sequences (figure 2), but for some of the species, the 5S arrangement has yet to be determined. Hence, we analyzed the 5S arrangement in all Asteraceae species and the three chosen outgroups for figure 2. This was done by low coverage read assembly, where the 35S and 5S contigs were visualized (figure 3). Arrangement-wise, we observed 10 separate and 5 linked rDNA arrangements within our dataset.

**Figure 3:**
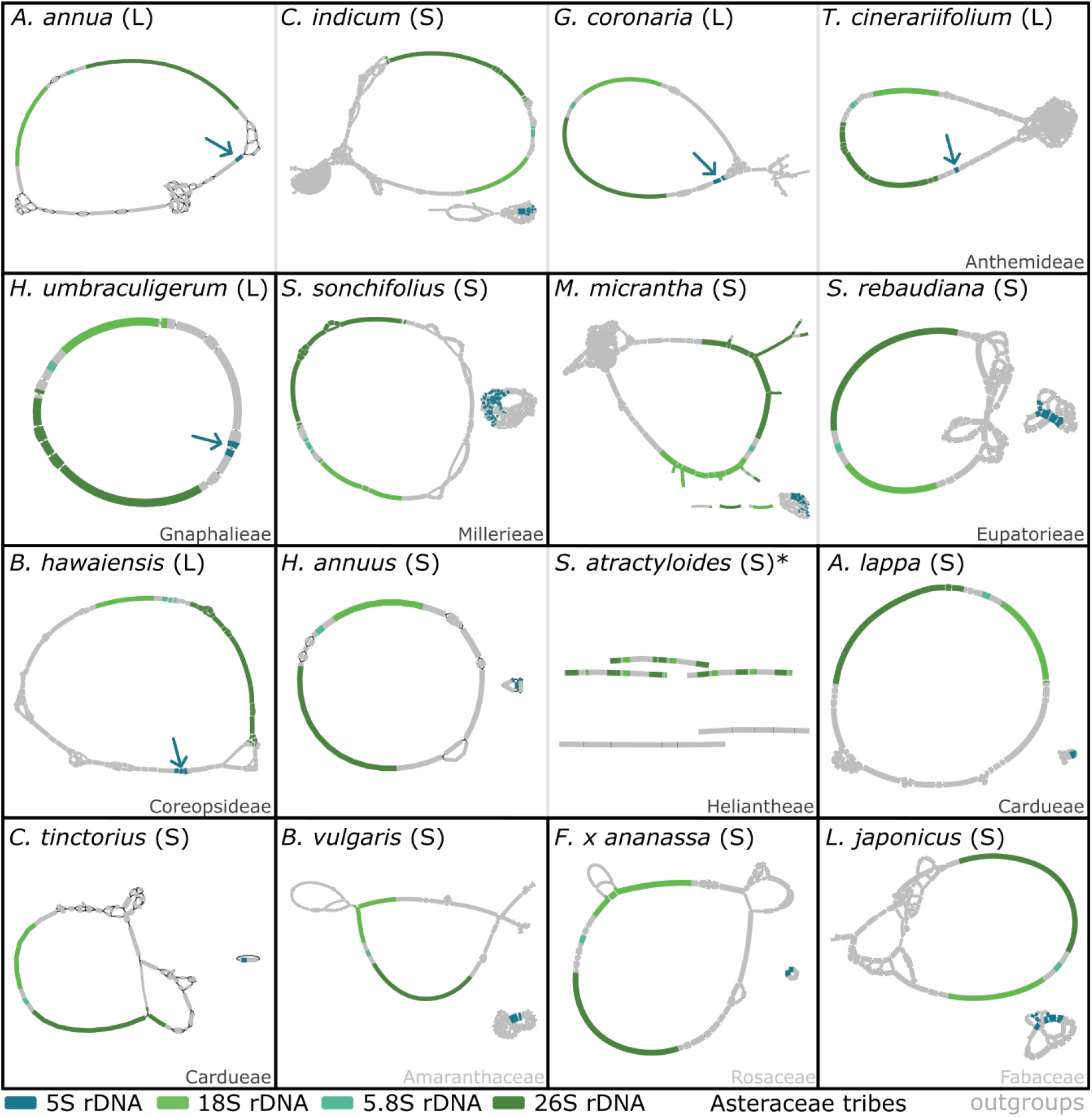
35S-5S linkage of ribosomal RNA genes is present in several, but not all, Asteraceae species. The graphs represent the low coverage assembly of the rRNA genes and show gene order, arrangement and organization. Complete circular graphs represent the typical rDNA monomer for each species. Linked ribosomal genes (L) show a single circular graph including the 5S rDNA gene (marked by arrows; five instances). Separated ribosomal RNA genes (S) show two graphs (10 instances). For *Scalesia atractyloides* (*), read quality strongly impaired the alignments and prevented full circular assemblies. Nevertheless, the rRNA genes were assembled completely, with several copies being separated by spacers. A separated arrangement can be concluded even from the imperfect assembly of *S. atractyloides* rDNAs. Five of the Asteraceae show the 35S-5S linkage being neither restricted to species with the promoter shift (see *Helichrysum umbraculigerum* and *Bidens hawaiensis*) nor affecting all species with the shifted promoter (see *Chrysanthemum indicum*). The line thickness is associated to the coverage depth of individual contigs; however, these are not comparable across the species shown. Sequence representation is not to scale.

We conclude that the Asteraceae species *Artemisia annua*, *Glebionis coronaria*, *Tanacetum cinerariifolium* (all within the Anthemideae), *Helichrysum umbraculigerum* (Gnaphalieae) and *Bidens hawaiensis* (Coreopsideae) have 35S-5S linkage. Clearly, despite 35S-5S linkage occurring often in the Anthemideae as reported, it is neither restricted to this tribe nor present in all species of this tribe. Instead, linkage occurs polyphyletically and is not necessarily associated with the promoter shift limited to the Anthemideae (figure 2). Nevertheless, considering linkage in the Anthemideae and its sister tribe Gnaphalieae, we assume that the linked 35S-5S arrangement emerged first and that the promoter sequences shifted later in the evolutionary timeline.

### Fixed in the 5S gene, but variable in the TE: Cassandra 5S regions show two variable sequence motifs

Having clarified that the promoter shifts and 35S-5S linkage are not always correlated, we wondered about the sequence variability between the promoter box motifs. If some motifs are more variable than others, they may be used to gain further insight in the evolutionary history of the 5S rRNA gene and the accompanying Cassandra retrotransposons: As highlighted in figure 2, the LTR core unit includes the highly conserved 5S rDNA promoter box motifs and parts of their flanking genic region. However, in Cassandra, not all parts of the similarity region are as highly conserved as the promoter box motifs.

Comparing Cassandra retrotransposon sequences to the 5S rRNA genes, we note that Cassandra retrotransposons have regions of high variability just next to the box motifs: In the promoter-flanking regions, an accumulation of mutations appears to be tolerated between the boxes (figure 4). The nucleotide sequence most prone to mutations thereby seems to be right in front of the conserved box motifs, e.g., between the A-Box and the Intermediate Element (IE), we observe 4 nt long and between the IE and the C-Box 5 nt long polymorphic motifs, further referred as MotIE and MotC (figure 4).

**Figure 4:**
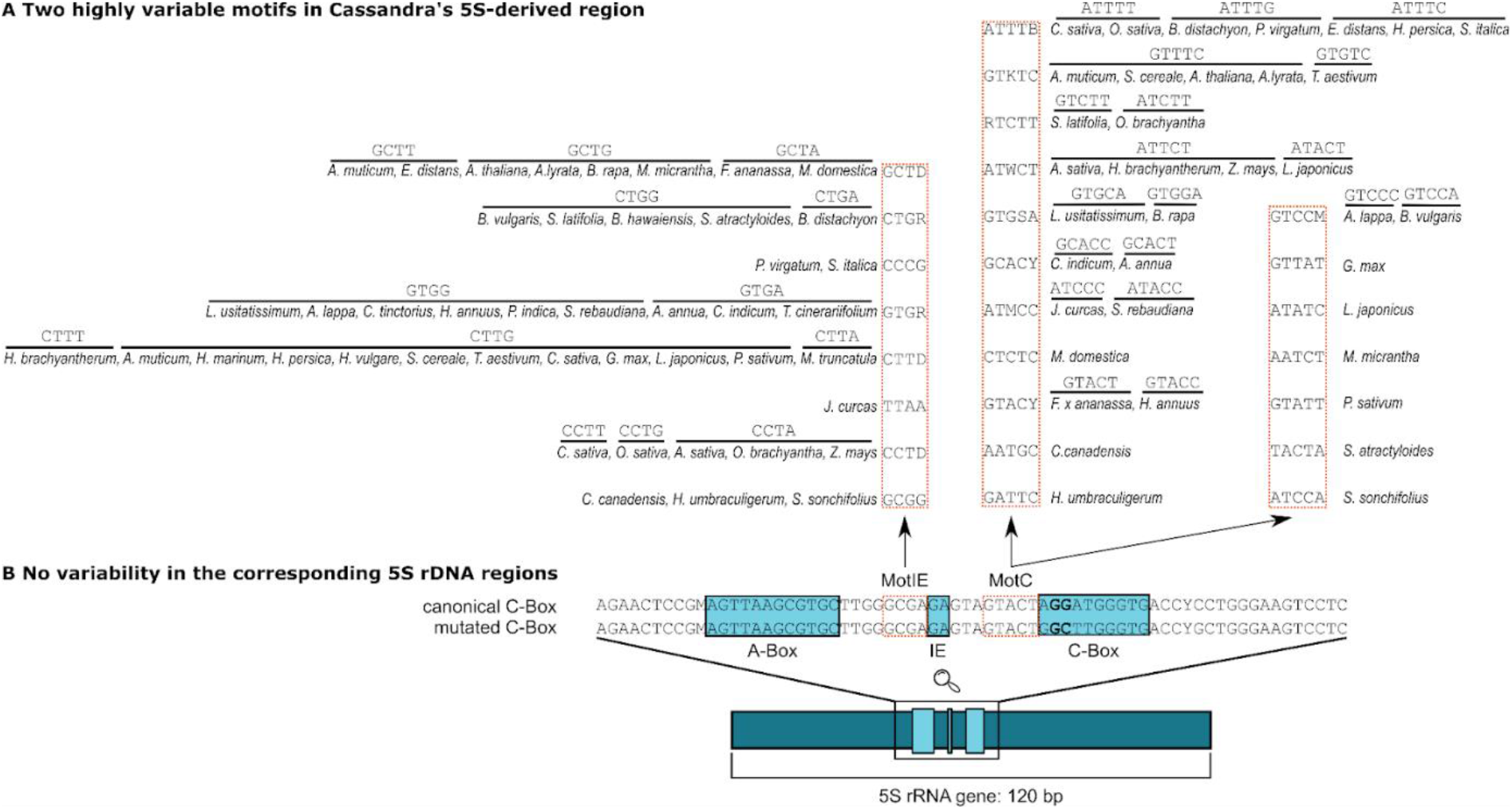
Polymorphic motifs in Cassandra LTR - 5S similarity region. Within the conserved 5S region within the Cassandra LTR we observe variable motifs between the highly conserved promoter boxes (A-Box, Intermediate Element - IE, C-Box). Depending on their localization and association with the nearest conserved promoter boxes, they are called MotIE and MotC. For MotIE and MotC we observe shifts in sequence information within the Cassandra LTRs (A). In contrast, there is no observable variability in the corresponding 5S rDNA genes in these regions (B). For 5S rDNA of corresponding genes consensuses for mutated and canonical C-Boxes are shown.

To better understand the regions of variability between the Cassandra LTRs, we further investigated the nucleotide compositions in the MotIE and MotC regions and how they map back to the 5S rDNA. Most striking are the differences in nucleotide composition of MotIE and MotC polymorphisms. 45 Cassandra sequences show a MotIE polymorphism with eight observable nucleotide motif variants (figure 4) and only one being species-specific (*J. curca*s). MotC divergence from the 5S rRNA gene was detectable in 37 out of 45 Cassandra sequences Here, variability is even higher and we observe 18 different motifs with nine of them being specific for species of Rosaceae (*M. domestica*), Fabaceae (*G. max*, *L. japonicus*, *P. sativum*) and Asteraceae (*C. canadensis*, *H. umbraculigerum*, *M. micrantha, S. atractyloides, S. sonchifolius*). Nevertheless, most motif changes appear as stochastic variation and carry no clear phylogenetic signal.

### Assembling a framework to analyze Cassandra and 5S rDNA co-evolution

To better understand the co-evolution mechanisms that accompany the interplay of Cassandra retrotransposons and the 5S rDNA, we collected comprehensive Cassandra data across the vascular plants. We focused especially on the Asteraceae, combining Cassandra information, 5S rDNA promoters and 35S-5S linkage to illustrate the variable landscape of the 5S-Cassandra interaction (figure 5). For Asteraceae we gathered data from 11 plant tribes and 22 plant families (figure 5; column 1). Limitations are in the data foundation, as for some species not all required sequencing reads were available (figure 5; column 2). Nevertheless, our data provides a comprehensive framework that – for the first time – allows tracing Cassandra evolution (figure 5; column 3) in comparison to 5S evolutionary changes, such as promoter and arrangement shifts (figure 5; column 4).

**Figure 5:**
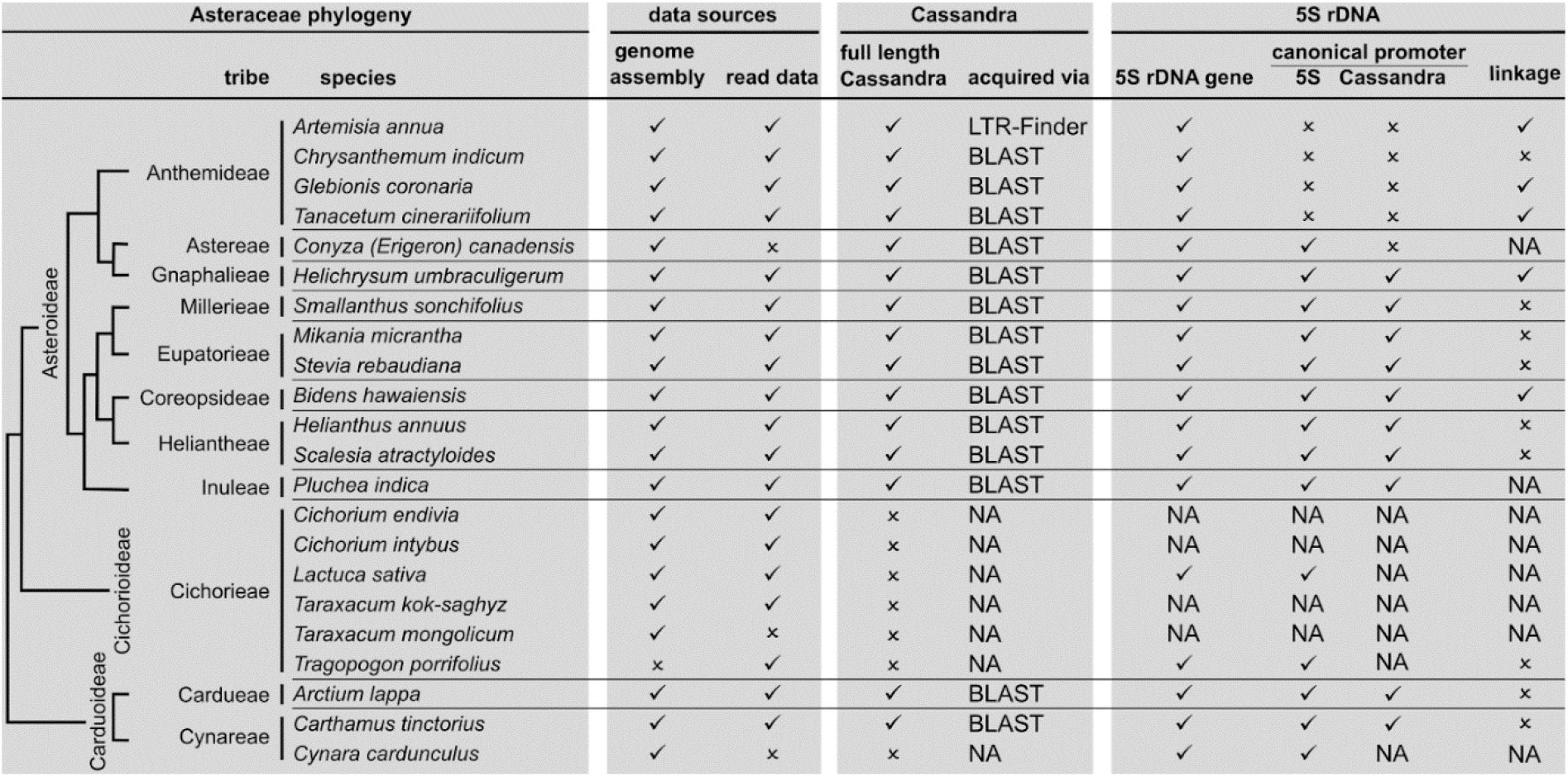
Framework for understanding the interplay between Cassandra retrotransposons and the 5S rDNA in the Asteraceae: Summary, integrative overview and data-based limitations. The Asteraceae phylogeny in the first column is based on Mandel et al. (2019).

## Discussion

### Plant Cassandras share structural hallmarks, but are variable in sequence and length: Cassandra is better defined as a lineage than as a family

As they are present across the plant kingdom, it could be assumed that Cassandra forms a single family of non-autonomous LTR retrotransposons. Our dataset of plant Cassandras now offers the most comprehensive framework up to date to re-examine, if all plant Cassandra sequences indeed form a single family. If we follow TE taxonomy, LTR retrotransposons are firstly classified into superfamilies and lineages according to their structure, e.g. by the presence and order of conserved regions, and in a second step by the sequence of the conserved regions (Wicker et al. 2007; Arkhipova 2017; Neumann et al. 2019). The main focus is usually on the enzymatic domains, as these are most conserved (Malik and Eickbush 1999; Malik 2005; Eickbush and Jamburuthugoda 2008). For family classification, LTR sequences are compared across sequences, but other parameters, such as element and LTR lengths as well as the internal sequences are also considered (Wicker et al. 2007). As Cassandra retrotransposons do not code for any proteins, we can only take into account these length and sequence parameters.

Across the plant kingdom, represented here by 18 plant families, 81 Cassandra retrotransposons share the 5S promoter as a central motif in the LTR. However, apart from this, they show large variability in element length, structure and nucleotide information (figure 1, table 1, suppl. table 1). For the LTR as a family-defining hallmark (Simon and Zimmerly 2008), we recognized family-specific sequence and length similarities beyond the conserved 5S-derived similarity region. Nevertheless, for Asteraceae and Fabaceae sequences, we surprisingly observe more variability in the LTRs (table 1) than in the internal region. This is caused by indels in the LTRs of the Asteraceae and Fabaceae Cassandras that can be interpreted as tribe/subfamily-specific phylogenetic signals. We observe a link between Cassandra diversification and species richness in the Asteraceae and Fabaceae, as these two belong to the most species-rich plant families, with various subfamilies (Christenhusz and Byng 2016). Therefore, it is not surprising that we see this variation also in repetitive elements. Regarding internal regions, we observe mostly preferred size ranges and conservation within plant families (i.e., within the Rosaceae and Fabaceae). Despite this general trend, the internal Cassandra regions can sometimes vary within the same family as observed in the Amaranthaceae (see also Maiwald et al. 2021 for an in-depth report). Concluding, apart from the 5S promoter as shared structural hallmark, Cassandra sequences and lengths are variable and can form several Cassandra families. Taxonomically, we therefore understand Cassandra retrotransposons as a lineage of non-autonomous LTR retrotransposons rather than a family. This is in line with the lineage concepts of autonomous LTR retrotransposons that are based on structural similarities (Llorens et al. 2009; Du et al. 2010). For example, TEs of the chromovirus lineage have a chromodomain in the integrase region (Neumann et al. 2011; Weber et al. 2013) and TEs of the Retand/Ogre lineage have two instead of a single ribonuclease H region (Neumann et al. 2019). If we put our observations in the frameworks for TE classification (see also the TE Hub initiative; Elliott et al. 2021), we can extend the lineage concept towards non-autonomous retrotransposons, and suggest the concept of a Cassandra lineage, encompassing all non-autonomous LTR retrotransposons that contain a similar 5S promoter region.

### Co-evolution of Cassandra and the 5S rDNA

For all Cassandras in our dataset for which we could retrieve the corresponding 5S rDNA gene, we observe mirroring of the conserved promoter box motifs. This is particularly noticeable within the Asteraceae, where different 5S variants and organizations have been described (Garcia et al. 2010; Garcia et al. 2012) and which we therefore investigated in detail (figure 5). We here describe Cassandra’s promoter mimicry in a depth that was not possible before. Nevertheless, promoter mimicry is not exclusively observed in Cassandra sequences. In plants, sequence mirroring strategies are widely found: “target mimicry”, for example, describes endogenous long non-coding RNAs that mimic and inhibit other small RNA molecules (miRNAs; Wu et al. 2013; Ye et al. 2014; Li et al. 2015). Although compared to these mechanisms, Cassandra promoter shifts may not affect genome integrity as strongly.

Regarding the mechanisms behind Cassandra promoter mimicry, an initial mutation within the 5S rDNA gene in an Anthemideae ancestor is a likely scenario. Due to concerted evolution of the ribosomal genes (Nei and Rooney 2005), mutations can spread through the array and become fixed. This is well in line with the current concerted evolution models of the 5S rDNA. We can assume, then, that a mutated C-Box in the 5S rDNA would be followed by molecular changes in the corresponding transcription factors on a cellular level to guarantee sufficient transcription.

As Cassandra retrotransposons rely on the 5S rDNA promoter for transcription (Kalendar et al. 2008; Maiwald et al. 2021), they need to either adapt or become inactive relics. This would also increase the mutation pressure for Cassandra sequences. Cassandra’s promoter mimicry that mirrors the mutation in the C-Box (either random or also caused by gene conversion) was likely necessary to maintain TE integrity and to avoid vanishing. As an alternative hypothesis, an independent emergence of the C-Box in Cassandra sequences within the Anthemideae spawned from each 5S rDNA variant could also be possible, but seems unlikely, as overall sequence identity between Anthemideae Cassandras and the other Asteraceae is too high.

We thus suggest the following chain of events: First, the rDNA promoter mutated, followed by homogenization/concerted evolution. Second, Cassandra retrotransposons mirrored these C-Box shifts to survive. Third, mobilization (and with this reverse transcription, amplification and integration) of Cassandra retrotransposons led to further copies with the new promoter. At the same time, spread through already existing Cassandra copies may have been influenced by gene conversion and even by homogenization in tandemly arranged Cassandras (as seen in Yin et al. 2014; Kalendar et al. 2020; Maiwald et al. 2021).

In contrast to the highly conserved 5S promoter box motifs and the observed cases of promoter mimicry, we observed hypervariable motifs in the immediate vicinity of the highly conserved promoter boxes in the Cassandra LTRs. The hypervariable motifs MotIE and MotC of Cassandra are most striking in this context, i.e., within the expected most conserved region of the already highly conserved 5S rRNA gene. However, we assume the necessary preservation of the promoter boxes to enable transcription, whereas mutations in other regions are tolerated and can at least reach as much variability as the non-5S-derived sequence regions. Interestingly, mutations in these hypervariable motifs did not include larger indels. So, retention of certain lengths and spacing between the conserved promoter boxes seems to be mandatory, likely to enable the folding of secondary structures, as suggested previously for both 5S rDNA C-box variants (Garcia et al. 2012).

### Did Cassandra retrotransposons emerge multiple times?

Despite having rebuked an independent Cassandra emergence in tribe Anthemideae, the question holds: Is it likely that Cassandra emerged several times in the plant kingdom? For the purpose of this study, we consider Cassandra emergence as the process of obtaining a 5S promoter in the LTR of a retrotransposon. The acquisition of new, prominent, genic promoters such as the 5S, would enable a retrotransposon to pursue new strategies for proliferation. This is already well known for non-autonomous retrotransposons, e.g., for SINES in animals (Vassetzky and Kramerov 2013). Even apart from promoter structures, many instances of co-option of other sequence modules by TEs were observed (reviewed by Cosby et al. 2019; Wang and Han 2021). We hypothesize that the emergence of Cassandra retrotransposons was caused by spatial association of an LTR retrotransposon near an existing or extrachromosomally located 5S rDNA array (figure 6A). Also, a direct integration of Cassandra into a 5S rDNA cluster is imaginable as this would enable a first quick Cassandra outburst through accessible regulatory transcription factors (Dewannieux et al. 2004; Lynch et al. 2015) before elimination due to 5S homogenization (Garcia et al., 2023). Either way, the close proximity of a Cassandra progenitor and nuclear 5S rDNA would have allowed promoter obtainment, without harming the 5S gene itself. An explanation for this neutral interaction could be a tendency to pseudogene accumulation at the borders of ribosomal RNA arrays (Robicheau et al. 2017).

**Figure 6:**
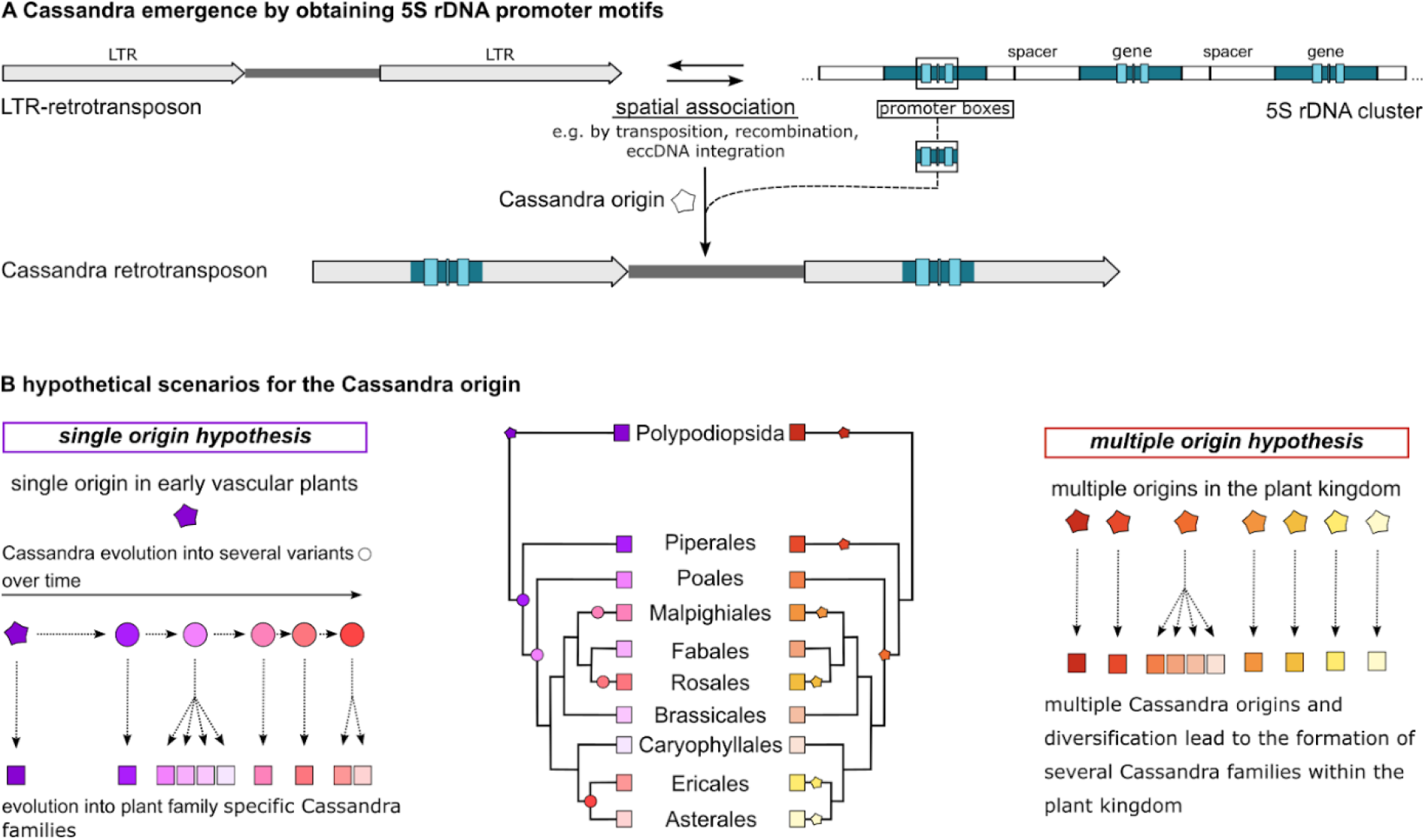
Cassandra origin and evolution within the plant kingdom. We suggest a Cassandra origin by obtaining 5S rDNA promoter (and flanking) regions due to spatial association, e.g., transposition, recombination or eccDNA integration through a LTR-retrotransposon (A). Based on our data we assume Cassandra originating (star) either as a single event in early vascular plants (B - left side; purple star) or multiple origins (B - right side; red/yellow stars) in the plant kingdom. In the single origin scenario one originator formed multiple variants (purple/pink circles). Each of the Cassandra families, we observe today (purple/pink rectangles), is a successor of one of these variants. For the multiple origin hypothesis, the scenario slightly changes with each family (red/yellow rectangles) being assignable to one of the newly emerged Cassandras (red/yellow stars) within the plant kingdom.

#### For Cassandra establishment we will discuss two hypotheses

On the one hand, Cassandra retrotransposons may have arisen only once in the early vascular plants (Figure 6B - left side), which led to a stable Cassandra population in ancestral species. The single origin hypothesis is supported by the strong conservation of the 5S promoter region – a 70 bp sequence stretch that extends well past the promoter boxes, as pointed previously. In this scenario, this 5S region was obtained by an LTR retrotransposon once and was retained through plant and Cassandra evolution. This retention across the evolution of vascular plants may well be possible. If TE populations do not harm or significantly alter gene expression, they tend to be tolerated by the host (Zhang and Mager 2012), which could have been the case for Cassandra 5S regions. Following this line of thought, the large sequence differences between Cassandra retrotransposons of different plant families arose as result of mutation and sequence reshuffling.

On the other hand, Cassandra retrotransposons may have originated at multiple time points during plant evolution (Figure 6B - right side), forming several independent Cassandra populations in the plant kingdom. If Cassandra emerged independently, we would consider the 70 bp region including the 5S promoter as a sequence module that was taken up multiple times by non-autonomous LTR retrotransposons, thereby forming Cassandra sequences repeatedly, in independent manners. Modular evolution is one of the typical strategies of TE evolution, occurring across all major TE clades and lineages (Smyshlyaev et al. 2013; Heitkam et al. 2014; Seibt et al. 2020).

In the light of both scenarios, the newly identified Cassandra-like retrotransposon – a TE with high similarity to Cassandra, but without the 5S promoter – could be a possible Cassandra precursor. By acquiring the 5S promoter module, a full Cassandra may have been formed from this Cassandra-like TRIM. Alternatively, these Cassandra-like elements may have arisen from Cassandras that have lost the 5S rDNA promoter. Nevertheless, losing such a beneficial hallmark doesn’t seem to be advantageous. Either way, after Cassandra emergence, a range of different evolutionary mechanisms act on these TEs, including the accumulation of mutations, rearrangements, recombination, reshuffling, and even polymerization. These processes caused Cassandra diversification into different variants which, through further diversification, formed the Cassandra families we observe today (figure 6B - both sides). Both scenarios can explain Cassandra retrotransposons with large LTR similarity within plant families (figure 1). Diversification of transposable elements due to read-through transcription (Xiao et al. 2008; Elrouby and Bureau 2010; Jiang and Ramachandran 2013), recombination (Devos et al. 2002; El Baidouri and Panaud 2013; Yin et al. 2014; Yin et al. 2015; Kalendar et al. 2020; Maiwald et al. 2021) and chimeric TE formation (Vicient et al. 2005; Wollrab et al. 2012; Sanchez et al. 2017) is a common mechanism. Also, the formation of extrachromosomal DNA during reverse transcription can influence TE sequence and structure on a nucleotide level (Drost and Sanchez 2019).

Whatever form of evolution has taken place, the 5S promoter is a powerful sequence module to be shuffled around, as it grants access to a reliable, environmental stress-independent transcription. The procurement of additional sequence information through reshuffling is known as a great step in LTR retrotransposon evolution (Malik and Eickbush 1999; Malik and Eickbush 2001) and, a part from conserved protein domains, this could also apply to promoter regions, as seen for SINEs (Seibt et al. 2020). The neat observed borders of the similarity region within all Cassandra LTRs (figure 2) clearly hint to some sort of modular reshuffling, with either the flanking LTR regions diversifying over time (single origin) or modular obtainment of this region by multiple “initiator” elements (multiple origin). Nevertheless, the strong similarity across the whole 5S region leads us to favor the single origin hypothesis.

### Has Cassandra carried the 5S gene into the linked arrangement?

35S-5S linkage is one the of the most prominent hallmarks of 5S rDNA in the Asteraceae. It is reported for several species (Garcia et al. 2010), but as a big controversy not necessarily in line with the recent phylogeny (Mandel et al. 2019). Our integrative dataset includes five Asteraceae with 5S rDNAs in linked arrangements (figure 5). We find that linkage is present in the Anthemideae, Gnaphalieae and Coreopsideae tribes, but in a patchy manner: First, phylogenetically, these three tribes are not direct sisters. Second, there is variation even within a tribe, as for instance, the Anthemideae contain both arrangement types (figure 3, figure 5; Garcia et al. 2009; Garcia et al. 2010). This patchy pattern of linked and separate arrangements across the phylogeny of the Asteraceae can be explained by either (1) multiple emergences of 35S-5S linkage or (2) a singular emergence of linkage followed by subsequent losses.

Regarding the multiple emergences of 35S-5S linkage in closely related Asteraceae, we consider this as unlikely. Despite reports of independent emergence for example in Ginkgo and other gymnosperm taxa (Garcia and Kovařík 2013), we consider the affected Asteraceae tribes as too closely related to have linkage occurring by chance.

Instead, we favor a single emergence, followed by loss of linkage within certain tribes/species and the retention in others. This scenario could be the result of frequent hybridization and polyploidization among the Asteraceae: If species with separate (S) and linked (L) arrangements hybridize, they would generate F1 hybrids (S×L) that have both arrangement types. One of them likely vanishes during the following generations (probably through concerted evolution mechanisms), thus enabling alternating and patchy distributions of S and L arrangements in and across populations. This scenario is supported by reports of frequent hybridization, polyploidization and general species richness in the Asteraceae (Semple and Watanabe 2023). Nevertheless, in both scenarios, there had to be a coexistence of linked and separate arrangements for a certain timespan after emergence, which can be seen by accidental observation of unlinked 5S rDNA units in an otherwise L-type species, *Coreopsis major* (Garcia et al. 2010). Clearly, for all scenarios, one 5S arrangement was retained whereas the other had to vanish – either by chance or by selection. On a molecular level, the fixation of one 5S arrangement type can be achieved by homogenization and concerted evolution processes.

As discussed for the promoter shifts, we see that Cassandra clearly follows 5S rDNA evolution. Therefore, we asked if Cassandra could also impact 35S-5S linkage emergence. At first glance it seems obvious: due to their mobility and copy-and-paste mode of proliferation, Cassandras could have carried the 5S rRNA gene into the 35S rDNA array, as considered previously (Garcia et al. 2009; Garcia et al. 2010). However, if we have a closer look at Cassandra itself, this seems very unlikely for two main reasons:

1. None of the 81 species-specific Cassandras in our dataset (from 17 different plant families all across the plant kingdom) carried the whole 5S rDNA gene. The region is always limited to a module containing the promoter regions and a short part of the flanking genic region.
2. Retrotransposons have a totally different structure compared to ribosomal DNA clusters, with other mechanisms putting evolutionary pressure on the sequence.

On the flip-side, we assume that 35S-5S linkage might have had an influence on Cassandra emergence and evolution. One can assume a limitation of 5S diversification through linkage, as concerted evolution may have a greater impact on 5S in linked arrangements as opposed to 5S in separated arrangements. Mutation rates support this hypothesis (Sònia Garcia et al., personal communication). If the linked arrangements allowed a faster spreading and fixation of 5S mutations in the array, it would be the perfect starting ground for the observed promoter shifts in the Anthemideae. As discussed above, in order to survive, Cassandra sequences then must adapt to this shift and mimic the new C-Box variant/information.

## Conclusion

We collated a dataset of 81 species-specific plant Cassandra retrotransposons and defined the presence of the 5S-related sequence stretch in the LTR as a hallmark defining the Cassandra TE lineage. Narrowing down on the Asteraceae, a plant family with wide variation in the 5S gene sequence and organization, we put together a comprehensive Cassandra-5S rDNA framework to trace the interplay between Cassandra TEs and 5S rDNA evolution. We find that shifts in 5S promoters are mimicked closely by the TE, whereas overall reorganization in 5S rDNA architecture does not impact the Cassandra landscapes. We here provide convincing evidence for gradual Cassandra-5S rDNA co-evolution that gives insight into the interplay between TEs and rDNA in plant genomes.

## Declarations

### Ethics approval and consent to participate

Not applicable

### Consent for publication

All authors have read and approved this manuscript.

### Availability of data and materials

Link to external archive

## Authors’ contributions

Analysis: SM, SG, LM

Writing: all

Figures: SM, LM

Conception: all

## Supporting information

Supplemental Table 1: Data mining

Supplemental Table 2: Cassandra and 5S metadata

## Acknowledgements

Open Access funding was enabled and organized by the Sächsische Staats-und Universitätsbibliothek and the Technische Universität Dresden. Daniel Vitales and Joan Pere Pascual-Díaz are acknowledged for help with 5S rDNA analysis. Computational resources were provided by the ELIXIR-CZ project (LM2015047), part of the international ELIXIR infrastructure.” (see https://repeatexplorer-elixir.cerit-sc.cz/)

## References

Alexandrov OS, Razumova OV, Karlov GI. 2021. A comparative study of 5S rDNA non-transcribed spacers in Elaeagnaceae species. Plants 10:4.

Andrews S. 2010. FASTQC. A quality control tool for high throughput sequence data. https://www.bioinformatics.babraham.ac.uk/projects/fastqc/.

Antonius-Klemola K, Kalendar R, Schulman AH. 2006. TRIM retrotransposons occur in apple and are polymorphic between varieties but not sports. Theor Appl Genet 112:999–1008.

Arkhipova IR. 2017. Using bioinformatic and phylogenetic approaches to classify transposable elements and understand their complex evolutionary histories. Mobile DNA 8:19.

Badouin H, Gouzy J, Grassa CJ, Murat F, Staton SE, Cottret L, Lelandais-Brière C, Owens GL, Carrère S, Mayjonade B, et al. 2017. The sunflower genome provides insights into oil metabolism, flowering and Asterid evolution. Nature 546:148–152.

Bankevich A, Nurk S, Antipov D, Gurevich AA, Dvorkin M, Kulikov AS, Lesin VM, Nikolenko SI, Pham S, Prjibelski AD, et al. 2012. SPAdes: A new genome assembly algorithm and its applications to single-cell sequencing. J Comput Biol 19:455–477.

Baum B, Johnson D. 2011. A comparison of the 5S rDNA diversity in the *Hordeum brachyantherum* californicum complex with those of the eastern Asiatic *Hordeum roshevitzii* and the South American *Hordeum cordobense* (Triticeae: Poaceae). Canadian Journal of Botany 80:752–762.

Baum BR, Bailey LG, Belyayev A, Raskina O, Nevo E. 2004. The utility of the nontranscribed spacer of 5S rDNA units grouped into unit classes assigned to haplomes - a test on cultivated wheat and wheat progenitors. Genome 47:590–599.

Baum BR, Edwards T, Johnson DA. 2013. What does the 5S rRNA multigene family tell us about the origin of the annual Triticeae (Poaceae)? Genome 56:245–266.

Baum BR, Johnson DA. 1994. The molecular diversity of the 5S rRNA gene in barley (*Hordeum vulgare*). Genome 37:992–998.

Baum BR, Johnson DA. 1998. The 5S rRNA gene in sea barley (*Hordeum marinum* Hudson sensu lato): sequence variation among repeat units and relationship to the X haplome in barley (*Hordeum*). Genome 41:652–661.

Bolger AM, Lohse M, Usadel B. 2014. Trimmomatic: a flexible trimmer for Illumina sequence data. Bioinformatics 30:2114–2120.

Bourque G, Burns KH, Gehring M, Gorbunova V, Seluanov A, Hammell M, Imbeault M, Izsvák Z, Levin HL, Macfarlan TS, et al. 2018. Ten things you should know about transposable elements. Genome Biol 19:199.

Brown DD, Wensink PC, Jordan E. 1972. A comparison of the ribosomal DNA’s of *Xenopus laevis* and *Xenopus mulleri*: the evolution of tandem genes. J Mol Biol 63:57–73.

Christenhusz MJM, Byng JW. 2016. The number of known plants species in the world and its annual increase. Phytotaxa 261:201.

Cohen S, Agmon N, Sobol O, Segal D. 2010. Extrachromosomal circles of satellite repeats and 5S ribosomal DNA in human cells. Mobile DNA 1:11.

Cosby RL, Chang N-C, Feschotte C. 2019. Host–transposon interactions: conflict, cooperation, and cooption. Genes Dev. 33:1098–1116.

Cullings K w. 1992. Design and testing of a plant-specific PCR primer for ecological and evolutionary studies. Mol Ecol 1:233–240.

Devos KM, Brown JKM, Bennetzen JL. 2002. Genome size reduction through illegitimate recombination counteracts genome expansion in *Arabidopsis*. Genome Res 12:1075–1079.

Dewannieux M, Dupressoir A, Harper F, Pierron G, Heidmann T. 2004. Identification of autonomous IAP LTR retrotransposons mobile in mammalian cells. Nat Genet 36:534–539.

Dohm JC, Minoche AE, Holtgräwe D, Capella-Gutiérrez S, Zakrzewski F, Tafer H, Rupp O, Sörensen TR, Stracke R, Reinhardt R, et al. 2014. The genome of the recently domesticated crop plant sugar beet (*Beta vulgaris*). Nature 505:546–549.

Doyle JJ, Doyle JL eds. 1987. A rapid DNA isolation procedure for small quantities of fresh leaf tissue. Phytochemical Bulletin:11–15.

Drost H-G, Sanchez DH. 2019. Becoming a selfish clan: Recombination associated to reverse-transcription in LTR retrotransposons. Genome Biol Evol 11:3382–3392.

Drouin G, de Sá MM. 1995. The concerted evolution of 5S ribosomal genes linked to the repeat units of other multigene families. Mol Biol Evol 12:481–493.

Du J, Tian Z, Hans CS, Laten HM, Cannon SB, Jackson SA, Shoemaker RC, Ma J. 2010. Evolutionary conservation, diversity and specificity of LTR-retrotransposons in flowering plants: insights from genome-wide analysis and multi-specific comparison. Plant J 63:584–598.

Eaves LA, Gardner AJ, Fry RC. 2020. Chapter 2 - Tools for the assessment of epigenetic regulation. In: Fry RC, editor. Environmental Epigenetics in Toxicology and Public Health. Vol. 22. Translational Epigenetics. Academic Press. p. 33–64. Available from: https://www.sciencedirect.com/science/article/pii/B9780128199688000020

Edgar RC. 2004. MUSCLE: multiple sequence alignment with high accuracy and high throughput. Nucleic Acids Res 32:1792–1797.

Eickbush TH, Jamburuthugoda VK. 2008. The diversity of retrotransposons and the properties of their reverse transcriptases. Virus Res 134:221–234.

El Baidouri M, Panaud O. 2013. Comparative genomic paleontology across plant kingdom reveals the dynamics of TE-driven genome evolution. Genome Biol Evol 5:954–965.

Elliott TA, Heitkam T, Hubley R, Quesneville H, Suh A, Wheeler TJ, The TE Hub Consortium. 2021. TE Hub: A community-oriented space for sharing and connecting tools, data, resources, and methods for transposable element annotation. Mobile DNA 12:16.

Ellis TH, Lee D, Thomas CM, Simpson PR, Cleary WG, Newman MA, Burcham KW. 1988. 5S rRNA genes in *Pisum*: sequence, long range and chromosomal organization. Mol Gen Genet 214:333–342.

Elrouby N, Bureau TE. 2010. Bs1, a new chimeric gene formed by retrotransposon-mediated exon shuffling in maize. Plant Physiol 153:1413–1424.

Fan W, Wang S, Wang H, Wang A, Jiang F, Liu H, Zhao H, Xu D, Zhang Y. 2022. The genomes of chicory, endive, great burdock and yacon provide insights into Asteraceae palaeo-polyploidization history and plant inulin production. Mol Ecol Resour 22:3124–3140.

Fulnecek J, Matyásek R, Kovarík A. 2002. Distribution of 5-methylcytosine residues in 5S rRNA genes in *Arabidopsis thaliana* and *Secale cereale*. Mol Genet Genomics 268:510–517.

Gao D, Li Y, Kim KD, Abernathy B, Jackson SA. 2016. Landscape and evolutionary dynamics of terminal repeat retrotransposons in miniature in plant genomes. Genome Biol 17:7.

Garcia S, Crhák Khaitová L, Kovařík A. 2012. Expression of 5 S rRNA genes linked to 35 S rDNA in plants, their epigenetic modification and regulatory element divergence. BMC Plant Biol 12:95.

Garcia S, Kovařík A. 2013. Dancing together and separate again: gymnosperms exhibit frequent changes of fundamental 5S and 35S rRNA gene (rDNA) organisation. Heredity 111:23–33.

Garcia S, Lim KY, Chester M, Garnatje T, Pellicer J, Vallès J, Leitch AR, Kovařík A. 2009. Linkage of 35S and 5S rRNA genes in *Artemisia* (family Asteraceae): first evidence from angiosperms. Chromosoma 118:85–97.

Garcia S, Panero JL, Siroky J, Kovarik A. 2010. Repeated reunions and splits feature the highly dynamic evolution of 5S and 35S ribosomal RNA genes (rDNA) in the Asteraceae family. BMC Plant Biology:18.

Goldsbrough PB, Ellis TH, Lomonossoff GP. 1982. Sequence variation and methylation of the flax 5S RNA genes. Nucleic Acids Res 10:4501–4514.

He Z, Feng X, Chen Q, Li L, Li S, Han K, Guo Z, Wang J, Liu M, Shi C, et al. 2022. Evolution of coastal forests based on a full set of mangrove genomes. Nat Ecol Evol 6:738–749.

Heitkam T, Holtgräwe D, Dohm JC, Minoche AE, Himmelbauer H, Weisshaar B, Schmidt T. 2014. Profiling of extensively diversified plant LINEs reveals distinct plant-specific subclades. Plant J 79:385–397.

Hemleben V, Grierson D, Borisjuk N, Volkov RA, Kovarik A. 2021. Personal perspectives on plant ribosomal RNA genes research: From precursor-rRNA to molecular evolution. Front Plant Sci 12.

Hisano H, Tsujimura M, Yoshida H, Terachi T, Sato K. 2016. Mitochondrial genome sequences from wild and cultivated barley (*Hordeum vulgare*). BMC Genomics 17:824.

Isobe SN, Shirasawa K, Nagano S, Hirakawa H. 2018. Current status of octoploid strawberry (*Fragaria × ananassa*) genome study. In: Hytönen T, Graham J, Harrison R, editors. The Genomes of Rosaceous Berries and Their Wild Relatives. Compendium of Plant Genomes. Cham: Springer International Publishing. p. 129–137. Available from: https://doi.org/10.1007/978-3-319-76020-9_10

Jiang S-Y, Ramachandran S. 2013. Genome-wide survey and comparative analysis of LTR retrotransposons and their captured genes in rice and sorghum. PLoS One 8:e71118.

Kalendar R, Raskina O, Belyayev A, Schulman AH. 2020. Long tandem arrays of Cassandra retroelements and their role in genome dynamics in plants. IJMS 21:2931.

Kalendar R, Schulman AH. 2006. IRAP and REMAP for retrotransposon-based genotyping and fingerprinting. Nat Protoc 1:2478–2484.

Kalendar R, Tanskanen J, Chang W, Antonius K, Sela H, Peleg O, Schulman AH. 2008. Cassandra retrotransposons carry independently transcribed 5S RNA. PNAS 105:5833–5838.

Li D, Luo R, Liu C-M, Leung C-M, Ting H-F, Sadakane K, Yamashita H, Lam T-W. 2016. MEGAHIT v1.0: A fast and scalable metagenome assembler driven by advanced methodologies and community practices. Methods 102:3–11.

Li F, Wang W, Zhao N, Xiao B, Cao P, Wu X, Ye C, Shen E, Qiu J, Zhu Q-H, et al. 2015. Regulation of Nicotine biosynthesis by an endogenous target mimicry of microRNA in tobacco. Plant Physiol 169:1062–1071.

Li H, Jain C, Bufallo V, Murray K, Langhorst B, Klötzl F. 2013. Seqtk: a fast and lightweight tool for processing FASTA or FASTQ sequences. https://github.com/lh3/seqtk [Internet]. Available from: https://github.com/lh3/seqtk

Li J, Wan Q, Abbott RJ, Rao G-Y. 2013. Geographical distribution of cytotypes in the *Chrysanthemum indicum* complex as evidenced by ploidy level and genome-size variation. J Syst Evol 51:196– 204.

Lin T, Xu X, Du H, Fan X, Chen Q, Hai C, Zhou Z, Su X, Kou L, Gao Q, et al. 2022. Extensive sequence divergence between the reference genomes of *Taraxacum kok-saghyz* and *Taraxacum mongolicum*. Sci China Life Sci 65:515–528.

Liu B, Yan J, Li W, Yin L, Li P, Yu H, Xing L, Cai M, Wang Hengchao, Zhao M, et al. 2020. *Mikania micrantha* genome provides insights into the molecular mechanism of rapid growth. Nat Commun 11:340.

Llorens C, Muñoz-Pomer A, Bernad L, Botella H, Moya A. 2009. Network dynamics of eukaryotic LTR retroelements beyond phylogenetic trees. Biol Direct 4:41.

Lynch VJ, Nnamani MC, Kapusta A, Brayer K, Plaza SL, Mazur EC, Emera D, Sheikh SZ, Grützner F, Bauersachs S, et al. 2015. Ancient transposable elements transformed the uterine regulatory landscape and transcriptome during the evolution of mammalian pregnancy. Cell Rep 10:551– 561.

Maiwald S, Weber B, Seibt KM, Schmidt T, Heitkam T. 2021. The Cassandra retrotransposon landscape in sugar beet (*Beta vulgaris*) and related Amaranthaceae: recombination and re-shuffling lead to a high structural variability. Ann Bot 127:91–109.

Malik HS. 2005. Ribonuclease H evolution in retrotransposable elements. Cytogenet Genome Res 110:392–401.

Malik HS, Eickbush TH. 1999. Modular evolution of the integrase domain in the Ty3/Gypsy class of LTR retrotransposons. J Virol 73:5186–5190.

Malik HS, Eickbush TH. 2001. Phylogenetic analysis of ribonuclease H domains suggests a late, chimeric origin of LTR retrotransposable elements and retroviruses. Genome Res 11:1187–1197.

Mandel JR, Dikow RB, Siniscalchi CM, Thapa R, Watson LE, Funk VA. 2019. A fully resolved backbone phylogeny reveals numerous dispersals and explosive diversifications throughout the history of Asteraceae. PNAS 116:14083–14088.

Mun T, Bachmann A, Gupta V, Stougaard J, Andersen SU. 2016. Lotus Base: An integrated information portal for the model legume *Lotus japonicus*. Sci Rep 6:39447.

Nei M, Rooney AP. 2005. Concerted and birth-and-death evolution of multigene families. Annu. Rev. Genet. 39:121–152.

Neumann P, Navrátilová A, Koblížková A, Kejnovský E, Hřibová E, Hobza R, Widmer A, Doležel J, Macas J. 2011. Plant centromeric retrotransposons: a structural and cytogenetic perspective. Mobile DNA 2:4.

Neumann P, Novák P, Hoštáková N, Macas J. 2019. Systematic survey of plant LTR-retrotransposons elucidates phylogenetic relationships of their polyprotein domains and provides a reference for element classification. Mobile DNA 10:1.

Novák P, Neumann P, Macas J. 2020. Global analysis of repetitive DNA from unassembled sequence reads using RepeatExplorer2. Nat Protoc 15:3745–3776.

Novák P, Neumann P, Pech J, Steinhaisl J, Macas J. 2013. RepeatExplorer: a Galaxy-based web server for genome-wide characterization of eukaryotic repetitive elements from next-generation sequence reads. Bioinformatics 29:792–793.

O’Connor CM, Adams JU. 2010. Essentials of Cell Biology. Cambridge: NPG Education Available from: https://www.nature.com/scitable/ebooks/essentials-of-cell-biology-14749010/

O’Neill K, Pirro S. 2020. The complete genome sequence of *Stevia rebaudiana*, the Sweetleaf. Available from: https://f1000research.com/articles/9-751

Pedrosa A, Sandal N, Stougaard J, Schweizer D, Bachmair A. 2002. Chromosomal map of the model legume *Lotus japonicus*. Genetics 161:1661–1672.

Peng Y, Lai Z, Lane T, Nageswara-Rao M, Okada M, Jasieniuk M, O’Geen H, Kim RW, Sammons RD, Rieseberg LH, et al. 2014. De novo genome assembly of the economically important weed Horseweed using integrated data from multiple sequencing platforms. Plant Physiol 166:1241– 1254.

Perina A, Seoane D, González-Tizón AM, Rodríguez-Fariña F, Martínez-Lage A. 2011. Molecular organization and phylogenetic analysis of 5S rDNA in crustaceans of the genus *Pollicipes* reveal birth-and-death evolution and strong purifying selection. BMC Evol Biol 11:304.

Rebordinos L, Cross I, Merlo A. 2013. High evolutionary dynamism in 5S rDNA of fish: state of the art. Cytogenet Genome Res 141:103–113.

Rey-Baños R, Sáenz de Miera LE, García P, Pérez de la Vega M. 2017. Obtaining retrotransposon sequences, analysis of their genomic distribution and use of retrotransposon-derived genetic markers in lentil (*Lens culinaris* Medik.). Kalendar R, editor. PLoS ONE 12:e0176728.

Reyes-Chin-Wo S, Wang Z, Yang X, Kozik A, Arikit S, Song C, Xia L, Froenicke L, Lavelle DO, Truco M-J, et al. 2017. Genome assembly with in vitro proximity ligation data and whole-genome triplication in lettuce. Nat Commun 8:14953.

Robicheau BM, Susko E, Harrigan AM, Snyder M. 2017. Ribosomal RNA genes contribute to the formation of pseudogenes and junk DNA in the human genome. Genome Biol Evol 9:380–397.

Sanchez DH, Gaubert H, Drost H-G, Zabet NR, Paszkowski J. 2017. High-frequency recombination between members of an LTR retrotransposon family during transposition bursts. Nat Commun 8:1283.

Sardouei-Nasab S, Nemati Z, Mohammadi-Nejad G, Haghi R, Blattner FR. 2023. Phylogenomic investigation of safflower (*Carthamus tinctorius*) and related species using genotyping-by-sequencing (GBS). Sci Rep 13:6212.

Scaglione D, Reyes-Chin-Wo S, Acquadro A, Froenicke L, Portis E, Beitel C, Tirone M, Mauro R, Lo Monaco A, Mauromicale G, et al. 2016. The genome sequence of the outbreeding globe artichoke constructed de novo incorporating a phase-aware low-pass sequencing strategy of F1 progeny. Sci Rep 6:19427.

Schmidt T, Schwarzacher T, Heslop-Harrison JS. 1994. Physical mapping of rRNA genes by fluorescent in-situ hybridization and structural analysis of 5S rRNA genes and intergenic spacer sequences in sugar beet (Beta vulgaris). Theoret. Appl. Genetics 88:629–636.

Schnable PS, Ware D, Fulton RS, Stein JC, Wei F, Pasternak S, Liang C, Zhang J, Fulton L, Graves TA, et al. 2009. The B73 maize genome: Complexity, diversity, and dynamics. Science 326:1112–1115.

Seibt KM, Schmidt T, Heitkam T. 2020. The conserved 3′ Angio-domain defines a superfamily of short interspersed nuclear elements (SINEs) in higher plants. Plant J 101:681–699.

Semple J, Watanabe K. 2023. An overview to the index to chromosome numbers in Asteraceae database: Revisiting base chromosome numbers, polyploidy, descending dysploidy, and hybridization. In: Garcia S, Nualart Deuxeus N, editors. Plant Genomics and Cytogenetic Databases. Vol. 2073. Methods in Molecular Biology.

Shen Q, Zhang L, Liao Z, Wang S, Yan T, Shi P, Liu M, Fu X, Pan Q, Wang Y, et al. 2018. The genome of *Artemisia annua* provides insight into the evolution of Asteraceae family and Artemisinin biosynthesis. Mol Plant 11:776–788.

Simon DM, Zimmerly S. 2008. A diversity of uncharacterized reverse transcriptases in bacteria. Nucleic Acids Res 36:7219–7229.

Smyshlyaev G, Voigt F, Blinov A, Barabas O, Novikova O. 2013. Acquisition of an Archaea-like ribonuclease H domain by plant L1 retrotransposons supports modular evolution. PNAS 110:20140–20145.

Song Y, Yang Y, Xu L, Bian C, Xing Y, Xue H, Hou W, Men W, Dou D, Kang T. 2023. The burdock database: a multi-omic database for *Arctium lappa*, a food and medicinal plant. BMC Plant Biology 23:86.

Staton SE, Bakken BH, Blackman BK, Chapman MA, Kane NC, Tang S, Ungerer MC, Knapp SJ, Rieseberg LH, Burke JM. 2012. The sunflower (*Helianthus annuus* L.) genome reflects a recent history of biased accumulation of transposable elements. Plant J 72:142–153.

Szymanski M, Zielezinski A, Barciszewski J, Erdmann VA, Karlowski WM. 2016. 5SRNAdb: an information resource for 5S ribosomal RNAs. Nucleic Acids Res 44:D180–D183.

The Angiosperm Phylogeny Group. 2017. APG IV: Angiosperm Phylogeny Group classification for the orders and families of flowering plants. Bot J Linn Soc 161:105–121.

Thibaud-Nissen F, DiCuccio M, Hlavina W, Kimchi A, Kitts PA, Murphy TD, Pruitt KD, Souvorov A. 2016. P8008 The NCBI eukaryotic genome annotation pipeline. Journal of Animal Science 94:184.

Vassetzky NS, Kramerov DA. 2013. SINEBase: a database and tool for SINE analysis. Nucleic Acids Res 41:D83–D89.

Vicient CM, Kalendar R, Schulman AH. 2005. Variability, recombination, and mosaic evolution of the barley BARE-1 retrotransposon. J Mol Evol 61:275–291.

Vierna J, Wehner S, Höner zu Siederdissen C, Martínez-Lage A, Marz M. 2013. Systematic analysis and evolution of 5S ribosomal DNA in metazoans. Heredity 111:410–421.

Waminal N, Ryu K, Park B, Kim H. 2014. Phylogeny of Cucurbitaceae species in Korea based on 5S rDNA non-transcribed spacer. Genes Genomics 36.

Wang J, Han G-Z. 2021. Unearthing LTR retrotransposon *gag* genes co-opted in the deep evolution of eukaryotes. Mol Biol Evol 38:3267–3278.

Wang S, Wang A, Wang H, Jiang F, Xu D, Fan W. 2022. Chromosome-level genome of a leaf vegetable *Glebionis coronaria* provides insights into the biosynthesis of monoterpenoids contributing to its special aroma. DNA Res 29:dsac036.

Weber B, Heitkam T, Holtgräwe D, Weisshaar B, Minoche AE, Dohm JC, Himmelbauer H, Schmidt T. 2013. Highly diverse chromoviruses of *Beta vulgaris* are classified by chromodomains and chromosomal integration. Mobile DNA 4:8.

Wick RR, Schultz MB, Zobel J, Holt KE. 2015. Bandage: interactive visualization of de novo genome assemblies. Bioinformatics 31:3350–3352.

Wicker T, Sabot F, Hua-Van A, Bennetzen JL, Capy P, Chalhoub B, Flavell A, Leroy P, Morgante M, Panaud O, et al. 2007. A unified classification system for eukaryotic transposable elements. Nat Rev Genet 8:973–982.

Witte C-P, Le QH, Bureau T, Kumar A. 2001. Terminal-repeat retrotransposons in miniature (TRIM) are involved in restructuring plant genomes. PNAS 98:13778–13783.

Wollrab C, Heitkam T, Holtgräwe D, Weisshaar B, Minoche AE, Dohm JC, Himmelbauer H, Schmidt T. 2012. Evolutionary reshuffling in the Errantivirus lineage Elbe within the *Beta vulgaris* genome. Plant J 72:636–651.

Wu H-J, Wang Z-M, Wang M, Wang X-J. 2013. Widespread long noncoding RNAs as endogenous target mimics for microRNAs in plants. Plant Physiol 161:1875–1884.

Wu Z, Liu H, Zhan W, Yu Z, Qin E, Liu S, Yang T, Xiang N, Kudrna D, Chen Y, et al. 2021. The chromosome-scale reference genome of safflower (*Carthamus tinctorius*) provides insights into linoleic acid and flavonoid biosynthesis. Plant Biotechnol J 19:1725–1742.

Xiao H, Jiang N, Schaffner E, Stockinger EJ, Van Der Knaap E. 2008. A retrotransposon-mediated gene duplication underlies morphological variation of Tomato fruit. Science 319:1527–1530.

Xu X, Yuan H, Yu X, Huang S, Sun Y, Zhang T, Liu Q, Tong H, Zhang Y, Wang Y, et al. 2021. The chromosome-level Stevia genome provides insights into steviol glycoside biosynthesis. Hortic Res 8:129.

Yamashiro T, Shiraishi A, Satake H, Nakayama K. 2019. Draft genome of *Tanacetum cinerariifolium*, the natural source of mosquito coil. Sci Rep 9:18249.

Ye C-Y, Xu H, Shen E, Liu Y, Wang Y, Shen Y, Qiu J, Zhu Q-H, Fan L. 2014. Genome-wide identification of non-coding RNAs interacted with microRNAs in soybean. Front Plant Sci 5.

Yin H, Du J, Li L, Jin C, Fan L, Li M, Wu J, Zhang S. 2014. Comparative genomic analysis reveals multiple long terminal repeats, lineage-specific amplification, and frequent interelement recombination for Cassandra retrotransposon in Pear (*Pyrus bretschneideri* Rehd.). Genome Biol Evol 6:1423–1436.

Yin H, Du J, Wu Jun, Wei S, Xu Y, Tao S, Wu Juyou, Zhang S. 2015. Genome-wide annotation and comparative analysis of long terminal repeat retrotransposons between Pear species of *P. bretschneideri* and *P. communis*. Sci Rep 5:17644.

Zhang Y, Mager DL. 2012. Gene properties and chromatin state influence the accumulation of transposable elements in genes. PLoS ONE 7:e30158.

Zhao M, Zhi H, Doust AN, Li W, Wang Yongfang, Li H, Jia G, Wang Yongqiang, Zhang N, Diao X. 2013. Novel genomes and genome constitutions identified by GISH and 5S rDNA and knotted1 genomic sequences in the genus *Setaria*. BMC Genomics 14:244.

