## Supplemental Figures 1-2 for "Evolving together: Cassandra retrotransposons gradually mirror promoter mutations of the 5S rRNA genes"

**Suppl. figure 1: Cassandra LTR alignment of different plant families.** Highlighting shows differences from consensus (A=red, T=green, C=yellow, G=blue). Consensus was calculated with a treshold of 25% (bases match atleast 25% of the sequences). For Fabaceae and Asteraceae Cassandras multiple Indels for certain species are observable leading to a split in different variants. Alignment data as fasta is provided on Zenodo (https://zenodo.org/record/8144620).

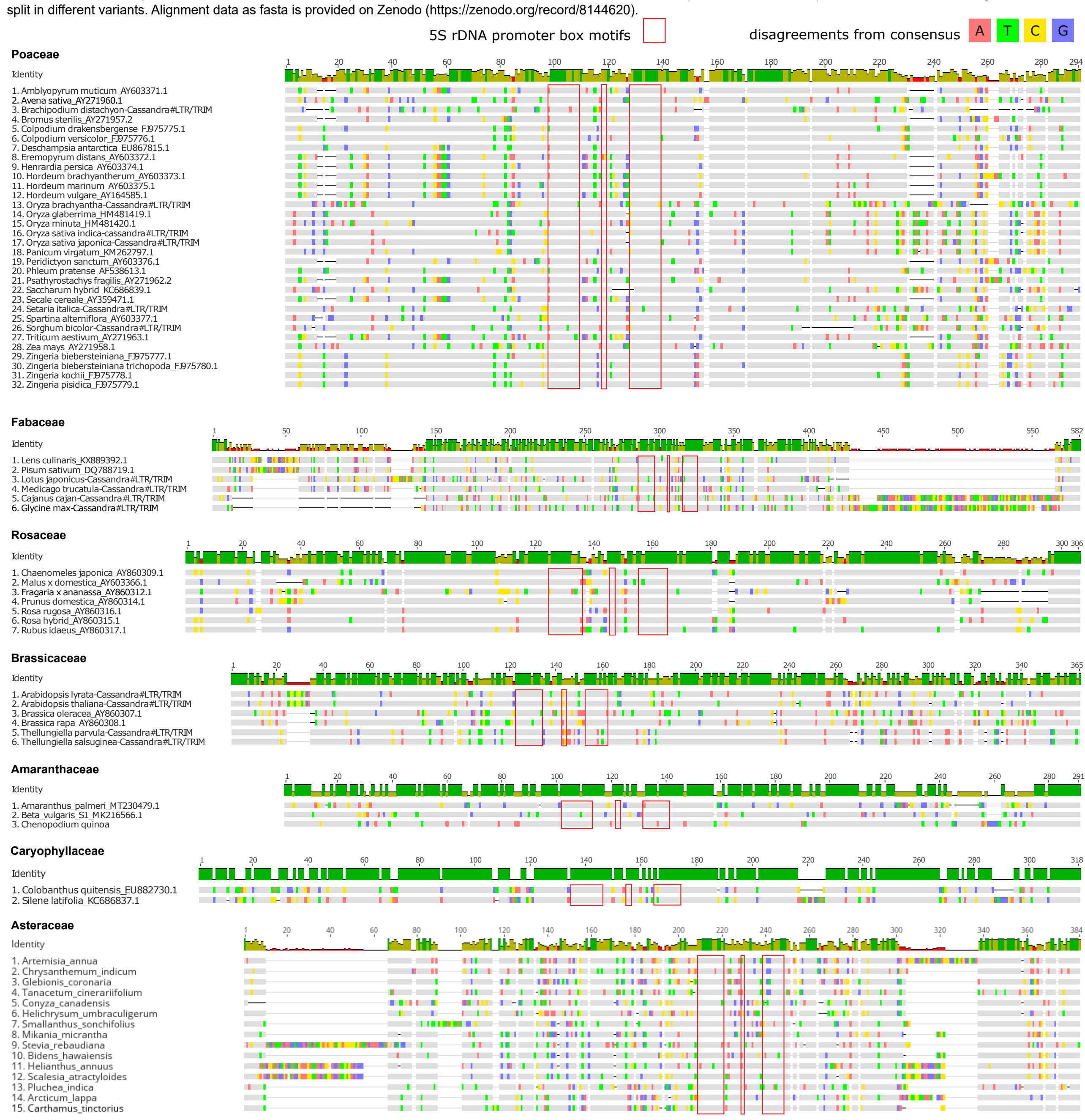


**Suppl. figure 2: Cassandra variants within the Asteraceae.** Two variants of Cassandra sequences can be identified within the Carduoideae. These sequences share the same internal region but show different LTR sequence information (A). LTR alignments of Asteraceae Cassandras show different LTR sequence informations for Cassandras and the Cassandra-like non autonomous elements (B). But internal regions of these variants are highly similiar, although Cassandra-like retrotransposons show a longer variant (C). As for *Bidens hawaiensis* (Bhaw) we observe a duplication within the internal region, leading to an usually longer variant. Differences from consensus are highlighted in color (A=red, T=green, C=yellow, G=blue). Consensus was calculated with a treshold of 25% (bases match atleast 25% of the sequences). Cassandra sequence names are shortened by species name: Aann = *Artemisia annua*, Cind = *Chrysanthemum indicum*, Gcor = *Glebionis coronaria*, Tcin = T*anacetum cinerariifolium*, Ccan = *Conyza canadensis*, Humb = *Helichrysum umbraculigerum*, Sson = *Smallanthus sonchifolius*, Mmic = *Mikania micrantha*, Sreb = *Stevia rebaudiana*, Bhaw = *Bidens hawaiensis*, Hann = *Helianthus annuus*, Satr = *Scalesia atractyloides*, Pind = *Pluchea indica*, Alap = *Arctium lappa*, Ctin = *Carthamus tinctorius*, Ccar = *Cynara cardunculus*

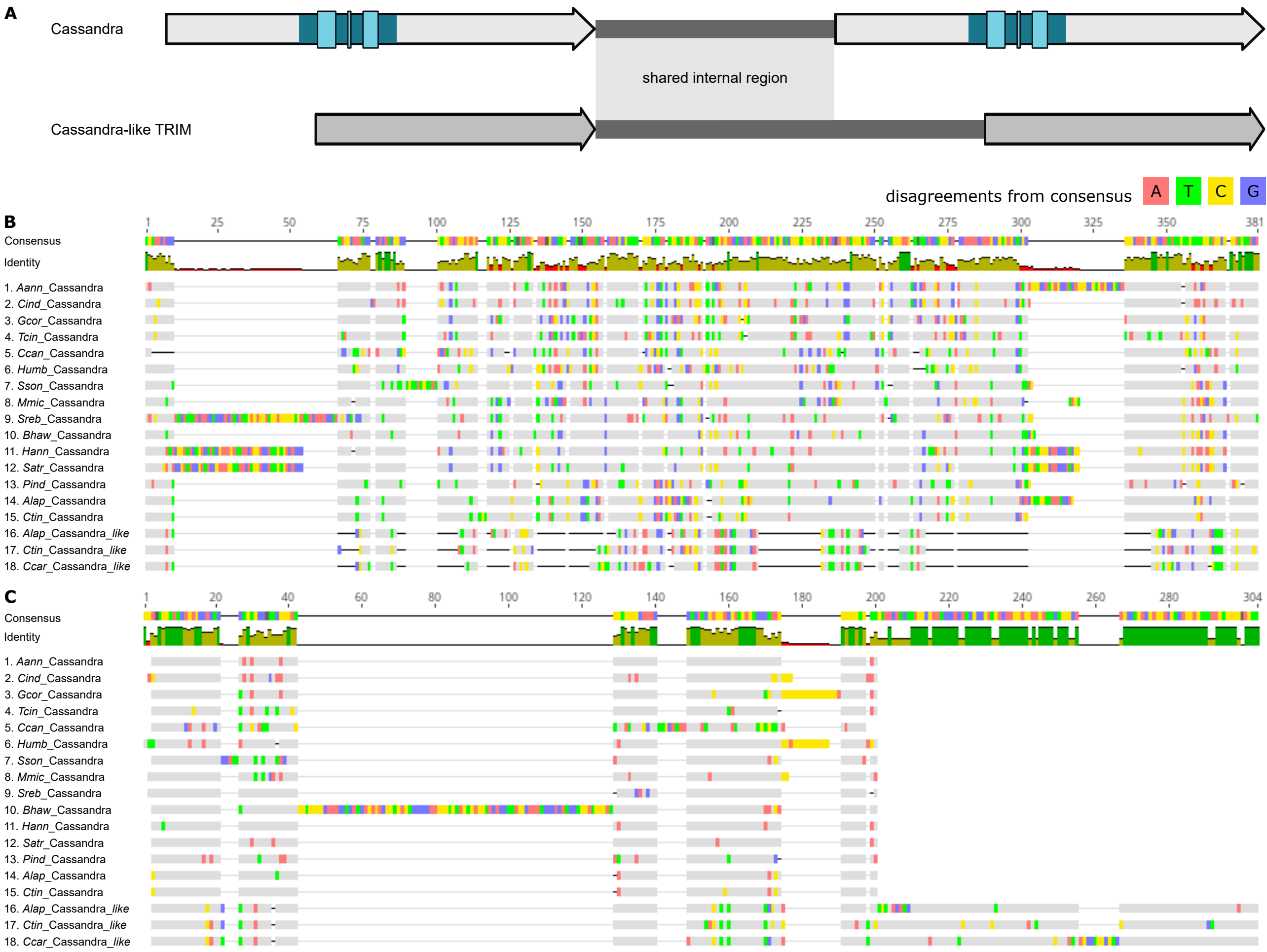
