## Supplemental Table 1: Data mining for "Evolving together: Cassandra retrotransposons gradually mirror promoter mutations of the 5S rRNA genes"

**Suppl. table 1A: Data sources for published Cassandra sequences**

| plant order | plant family | species | accession number | reference | DOI |
|---|---|---|---|---|---|
| Polypodiales | Didymochlaenaceae | *Didymochlaena truncutata* | AY860311.1 | Kalendar et al., 2008, | 10.1073/pnas.0709698105 |
| Polypodiales | Nephrolepidaceae | *Nephrolepis exaltata* | AY860313.1 | Kalendar et al., 2008 | 10.1073/pnas.0709698105 |
| Cyatheales | Cyatheaceae | *Sphaeropteris cooperi* | AY860310.1 | Kalendar et al., 2008 | 10.1073/pnas.0709698105 |
| Piperales | Aristolochiaceae | *Saruma henryi* | EF125873.1 | Kalendar et al., 2008 | 10.1073/pnas.0709698105 |
| Poales | Poaceae | *Ambylopyrum muticum* | AY603371.1 | Kalendar et al., 2008 | 10.1073/pnas.0709698105 |
| | Poaceae | *Avena sativa* | AY271960.1 | Kalendar et al., 2008 | 10.1073/pnas.0709698105 |
| | Poaceae | *Brachypodium distachyon* | Supplemental material from reference | Gao et al., 2016 | 10.1186/s13059-015-0867-y |
| | Poaceae | *Bromus sterilis* | AY271957 | Kalendar et al., 2008 | 10.1073/pnas.0709698105 |
| | Poaceae | *Colpodium drakensbergense* | FJ975775.1 | Kalendar et al., 2020 | 10.3390/ijms21082931 |
| | Poaceae | *Colpodium versicolor* | FJ975776.1 | Kalendar et al., 2020 | 10.3390/ijms21082931 |
| | Poaceae | *Deschampsia antarctica* | EU867815 | Kalendar et al., 2020 | 10.3390/ijms21082931 |
| | Poaceae | *Eremopyrum distans* | AY603372.1 | Kalendar et al., 2008 | 10.1073/pnas.0709698105 |
| | Poaceae | *Henrardia persica* | AY603374.1 | Kalendar et al., 2008 | 10.1073/pnas.0709698105 |
| | Poaceae | *Hordeum brachyantherum* | AY603373.1 | Kalendar et al., 2008 | 10.1073/pnas.0709698105 |
| | Poaceae | *Hordeum marinum* | AY603375.1 | Kalendar et al., 2008 | 10.1073/pnas.0709698105 |
| | Poaceae | *Hordeum vulgare* | AY164585.1 | Kalendar et al., 2008 | 10.1073/pnas.0709698105 |
| | Poaceae | *Oryza brachyantha* | Supplemental material from reference | Gao et al., 2016 | 10.1186/s13059-015-0867-y |
| | Poaceae | *Oryza glaberrima* | HM481419.1 | Kalendar et al., 2020 | 10.3390/ijms21082931 |
| | Poaceae | *Oryza minuta* | HM481420.1 | Kalendar et al., 2020 | 10.3390/ijms21082931 |
| | Poaceae | *Oryza sativa indica* | Supplemental material from reference | Gao et al., 2016 | 10.1186/s13059-015-0867-y |
| | Poaceae | *Oryza sativa japonica* | Supplemental material from reference | Gao et al., 2016 | 10.1186/s13059-015-0867-y |
| | Poaceae | *Panicum virgatum* | KM262797.1 | Kalendar et al., 2020 | 10.3390/ijms21082931 |
| | Poaceae | *Peridictyon sanctum* | AY603376.1 | Kalendar et al., 2008 | 10.1073/pnas.0709698105 |
| | Poaceae | *Phleum pratense* | AF538613.1 | unpublished | |
| | Poaceae | *Psathyrostachys fragilis* | AY271962.2 | Kalendar et al., 2008 | 10.1073/pnas.0709698105 |
| | Poaceae | *Saccharum hybrid* | KC686839.1 | Kalendar et al., 2020 | 10.3390/ijms21082931 |
| | Poaceae | *Secale cereale* | AY359471.1 | Kalendar et al., 2008 | 10.1073/pnas.0709698105 |
| | Poaceae | *Setaria italica* | Supplemental material from reference | Gao et al., 2016 | 10.1186/s13059-015-0867-y |
| | Poaceae | *Spartina alterniflora* | AY603377.1 | Kalendar et al., 2008 | 10.1073/pnas.0709698105 |
| | Poaceae | *Sorghum bicolor* | Supplemental material from reference | Gao et al., 2016 | 10.1186/s13059-015-0867-y |

| | | | | | |
|---|---|---|---|---|---|
| | Poaceae | *Triticum aestivum* | AY271963.1 | Kalendar et al., 2008 | 10.1073/pnas.0709698105 |
| | Poaceae | *Zea mays* | AY271958.1 | Kalendar et al., 2008 | 10.1073/pnas.0709698105 |
| | Poaceae | *Zingeria biebersteiniana* ssp. *trichopoda* | FJ975780.1 | Kalendar et al. 2020 | 10.3390/ijms21082931 |
| | Poaceae | *Zingeria biebersteiniana* | FJ975777.1 | Kalendar et al. 2020 | 10.3390/ijms21082931 |
| | Poaceae | *Zingeria kochii* | FJ975778.1 | Kalendar et al. 2020 | 10.3390/ijms21082931 |
| | Poaceae | *Zingeria pisidica* | FJ975779.1 | Kalendar et al. 2020 | 10.3390/ijms21082931 |
| Malpighiales | Clusiaceae | *Garcinia mangostana* | EU140956.1 | Kalendar et al., 2020 | 10.3390/ijms21082931 |
| | Linaceae | *Linum usitatissimum* | DQ767972.1 | Kalendar et al., 2008 | 10.1073/pnas.0709698105 |
| | Euphorbiaceae | *Jatropha curcas* | Supplemental material from reference | Gao et al., 2016 | 10.1186/s13059-015-0867-y |
| Fabales | Fabaceae | *Cajanus cajan* | Supplemental material from reference | Gao et al., 2016 | 10.1186/s13059-015-0867-y |
| | Fabaceae | *Glycine max* | Supplemental material from reference | Gao et al., 2016 | 10.1186/s13059-015-0867-y |
| | Fabaceae | *Lens culinaris* | KX889392 | Rey-Banos et al., 2016, | 10.1371/journal.pone.0176728 |
| | Fabaceae | *Lotus japonicus* | Supplemental material from reference | Gao et al., 2016 | 10.1186/s13059-015-0867-y |
| | Fabaceae | *Medicago truncatula* | Supplemental material from reference | Gao et al., 2016 | 10.1186/s13059-015-0867-y |
| | Fabaceae | *Pisum sativum* | DQ788719.1 | Kalendar et al., 2008 | 10.1073/pnas.0709698105 |
| Rosales | Cannabaceae | *Cannabis sativa* | Supplemental material from reference | Gao et al., 2016 | 10.1186/s13059-015-0867-y |
| | Rosaceae | *Chaenomeles japonica* | AY860309.1 | Kalendar et al., 2008 | 10.1073/pnas.0709698105 |
| | Rosaceae | *Fragaria x ananassa* | AY860312.1 | Kalendar et al., 2008 | 10.1073/pnas.0709698105 |
| | Rosaceae | *Malus domestica* | AY603366.1 | Kalendar et al., 2008 | 10.1073/pnas.0709698105 |
| | Rosaceae | *Prunus domestica* | AY860314.1 | Kalendar et al., 2008 | 10.1073/pnas.0709698105 |
| | Rosaceae | *Rosa hybrid* | AY860315.1 | Kalendar et al., 2008 | 10.1073/pnas.0709698105 |
| | Rosaceae | *Rosa rugosa* | AY860316.1 | Kalendar et al., 2008 | 10.1073/pnas.0709698105 |
| | Rosaceae | *Rubus idaeus* | AY860317.1 | Kalendar et al., 2008 | 10.1073/pnas.0709698105 |
| Brassicales | Brassicaceae | *Arabidopsis lyrata* | Supplemental material from reference | Gao et al., 2016 | 10.1186/s13059-015-0867-y |
| | Brassicaceae | *Arabidopsis thaliana* | Supplemental material from reference | Gao et al., 2016 | 10.1186/s13059-015-0867-y |
| | Brassicaceae | *Brassica oleracea* | AY860307.1 | Kalendar et al., 2008 | 10.1073/pnas.0709698105 |
| | Brassicaceae | *Brassica rapa* | AY860308.1 | Kalendar et al., 2008 | 10.1073/pnas.0709698105 |
| | Brassicaceae | *Thellungiella parvula* | Supplemental material from reference | Gao et al., 2016 | 10.1186/s13059-015-0867-y |
| | Brassicaceae | *Thellungiella salsuginea* | Supplemental material from reference | Gao et al., 2016 | 10.1186/s13059-015-0867-y |
| Caryophyllales | Aioaceae | *Mesembryanthemum crystallinum* | AY603370.1 | Kalendar et al., 2008 | 10.1073/pnas.0709698105 |
| | Amaranthaceae | *Amaranthus palmeri* | MT230479.1 | Kalendar et al., 2020 | 10.3390/ijms21082931 |

| | | | | | |
|---|---|---|---|---|---|
| | Amaranthaceae | *Beta vulgaris* | MK216566.1 | Maiwald et al., 2021 | 10.1093/aob/mcaa176 |
| | Amaranthaceae | *Chenopodium quinoa* | – | Maiwald et al., 2021 | 10.1093/aob/mcaa176 |
| | Caryophyllaceae | *Colobanhus quitensis* | EU882730.1 | Kalendar et al., 2020 | 10.3390/ijms21082931 |
| | Caryophyllaceae | *Silene latifolia* | KC686837.1 | Kalendar et al., 2020 | 10.3390/ijms21082931 |
| Ericales | Ericaceae | *Vaccinium corymbosum* | DQ673669.1 | Kalendar et al., 2008 | 10.1073/pnas.0709698105 |

**Suppl. table 1B: Data sources for published Asteraceae genomes**

| plant family | lineage | species | accession number | platform | reference | DOI |
|---|---|---|---|---|---|---|
| Asteraceae | Asteroideae | *Artemisia annua* | PKPP00000000 | ENA | Shen *et al* ., 2018 | 10.1016/j.molp.2018.03.015 |
| | | *Bidens hawaiiensis* | GCA_021521975.1 | ENA | Bellinger *et al* ., 2022 | 10.1093/jhered/esab077 |
| | | *Chrysanthemum indicum* | GWHBHNH00000000 | genome warehouse | – | |
| | | *Conyza (Erigeron) canadensis* | JSWR01000000 | ENA | Peng *et al* ., 2014 | 10.1104/pp.114.247668 |
| | | *Glebionis cornaria* | JANFOE000000000.1 | ENA | Wang *et al* ., 2022 | 10.1093/dnares/dsac036 |
| | | *Helianthus anuus* | GCF_002127325.2 | ENA | Badouin *et al* ., 2017 | 10.1038/nature22380 |
| | | *Helichrysum umbraculigerum* | CATIUR010000000 | ENA | Berman *et al* ., 2023 | 10.1038/s41477-023-01402-3 |
| | | *Mikania micrantha* | GCA_009363875.1 | ENA | Liu *et al* ., 2020 | 10.1038/s41467-019-13926-4 |
| | | *Pluchea indica* | GWHBCJV0000000 | genome warehouse | He *et al* ., 2022 | 10.1038/s41559-022-01744-9 |
| | | *Scalesia atractyloides* | https://doi.org/10.5061/dryad.8gtht76rh | dryad | Cerca *et al* ., 2022 | 10.1038/s41467-022-31280-w |
| | | *Smallanthus sonchifolius* | JAKNSE010000000 | ENA | Fan *et al* ., 2022 | 10.1111/1755-0998.13675 |
| | | *Stevia rebaudiana* | GCA_009936405 | ENA | Xu *et al* ., 2021 | 10.1038/s41438-021-00565-4 |
| | | *Tanacetum cinerariifolium* | BKCJ000000000.1 | ENA | Yamashiro *et al* ., 2019 | 10.1038/s41598-019-54815-6 |
| | Cichorioideae | *Cichorium endivia* | GCA_023376185.1 | ENA | Fan *et al* ., 2022 | 10.1111/1755-0998.13675 |
| | | *Cichorium intybus* | JAKNSD000000000 | ENA | Fan *et al* ., 2022 | 10.1111/1755-0998.13675 |
| | | *Lactuca sativa* | GCF_002870075.4 | ENA | Reyes-Chin-Wo *et al* ., 2017 | 10.1038/ncomms14953 |
| | | *Taraxakum kok-saghyz* | GWHBCHF00000000 | genome warehouse | Lin *et al* ., 2021 | 10.1007/s11427-021-2033-2 |
| | | *Taraxacum mongolicum* | GWHBCHG00000000 | genome warehouse | Lin *et al* ., 2021 | 10.1007/s11427-021-2033-2 |
| | Carduoideae | *Arctium lappa* | GCA_023525745.1 | ENA | Fan *et al* ., 2022 | 10.1111/1755-0998.13675 |
| | | *Carthamus tinctorius* | GCA_001633085.1 | ENA | Wu *et al* ., 2021 | 10.1111/pbi.13586 |
| | | *Cynara cardunculus* | GCA_001531365.2 | ENA | Scaglione *et al* ., 2016 | 10.1038/srep19427 |

**Suppl. table 1C: Data sources for published 5S rRNA genes**

| species | accession number | repository | reference | DOI |
|---|---|---|---|---|
| *Amblyopyrum muticum* | EU924818 | NCBI | Baum et al., 2009 | 10.1139/g03-146 |
| *Arabidopsis lyrata* | E02158 | 5S rRNAdb | Szymanski et al., 2016 | 10.1093/nar/gkv1081 |
| *Arabidopsis thaliana* | E00006 | 5S rRNAdb | Szymanski et al., 2016 | 10.1093/nar/gkv1081 |
| *Avena sativa* | EF071696 | NCBI | Peng et al., 2008 | 10.1104/pp.114.247668 |
| *Beta vulgaris* | Z25804 | NCBI | Schmidt et al., 1994 | 10.1007/BF01253964 |
| *Brachypodium distachyon* | XR_002960580 | NCBI | Thibaud-Nissen et al., 2016 | 10.2527/jas2016.94supplement4184x |
| *Brassica rapa* | E02489 | 5S rRNAdb | Szymanski et al., 2016 | 10.1093/nar/gkv1081 |
| *Cannabis sativa* | XR_004008092 | NCBI | Thibaud-Nissen et al., 2016 | 10.2527/jas2016.94supplement4184x |
| *Cajanus cajan* | XR_003803880 | NCBI | Thibaud-Nissen et al., 2016 | 10.2527/jas2016.94supplement4184x |
| *Chrysanthemum indicum* | OK181863 | NCBI | - | |
| *Cynara cardunculus* | XR_003070488 | NCBI | Thibaud-Nissen et al., 2016 | 10.2527/jas2016.94supplement4184x |
| *Eremopyrum distans* | KC188473 | NCBI | Baum et al., 2013 | 10.1139/gen-2012-0195 |
| *Fragaria x ananassa* | E00852 | 5S rRNAdb | Szymanski et al., 2016 | 10.1093/nar/gkv1081 |
| *Glycine max* | XR_005890139 | NCBI | Thibaud-Nissen et al., 2016 | 10.2527/jas2016.94supplement4184x |
| *Henrardia persica* | KC188485 | NCBI | Baum et al., 2013 | 10.1139/gen-2012-0195 |
| *Hordeum brachyantherum* | AY034775 | NCBI | Baum & Johnson, 2011 | 10.1139/b02-057 |
| *Hordeum marinum* | AF027583 | NCBI | Baum & Johnson, 1998 | PMID: 9809436 |
| *Hordeum vulgare* | HVU07378 | NCBI | Baum & Johnson, 1994 | 10.1139/g94-140 |
| *Jatropha curcas* | E00420 | 5S rRNAdb | Szymanski et al., 2016 | 10.1093/nar/gkv1081 |
| *Lactuca sativa* | E00211 | 5S rRNAdb | Szymanski et al., 2016 | 10.1093/nar/gkv1081 |
| *Linum usitatissimum* | X01531 | NCBI | Goldsbrough et al., 1982 | 10.3390/ijms22031302 |
| *Lotus japonicus* | AY040715 | NCBI | Pedrosa et al., 2002 | 10.1093/genetics/161.4.1661 |
| *Malus domestica* | XR_003771672 | NCBI | Thibaud-Nissen et al., 2016 | 10.2527/jas2016.94supplement4184x |
| *Medicago truncatula* | XR_005644521 | NCBI | Thibaud-Nissen et al., 2016 | 10.2527/jas2016.94supplement4184x |
| *Oryza brachyantha* | XR_005812828 | NCBI | Thibaud-Nissen et al., 2016 | 10.2527/jas2016.94supplement4184x |
| *Oryza sativa* | E00234 | 5S rRNAdb | Szymanski et al., 2016 | 10.1093/nar/gkv1081 |
| *Panicum virgatum* | XR_005680461 | NCBI | Thibaud-Nissen et al., 2016 | 10.2527/jas2016.94supplement4184x |
| *Pisum sativum* | AY499178 | NCBI | Ellis et al., 1988 | 10.1007/BF00337732 |
| *Pyrus breitschneiderii* | E02495 | 5S rRNAdb | Szymanski et al., 2016 | 10.1093/nar/gkv1081 |
| *Secale cereale* | AJ307365 | NCBI | Fulnecek et al., 2002 | 10.1007/s00438-002-0761-7 |
| *Setaria italica* | KC525411 | NCBI | Zhao et al., 2013 | 10.1186/1471-2164-14-244 |
| *Silene latifolia* | AB027248 | NCBI | - | |
| *Triticum aestivum* | E00299 | 5S rRNAdb | Szymanski et al., 2016 | 10.1093/nar/gkv1081 |
| *Zea mays* | E00011 | 5S rRNAdb | Szymanski et al., 2016 | 10.1093/nar/gkv1081 |

### Suppl. table 1D: Data sources for used plant read data

**data for Asteraceae species**

| species | read accession | platform | library layout | library strategy | library source | read length |
|---|---|---|---|---|---|---|
| *Arctium lappa* | ERR5554584 | illumina HiSeq 2500 | paired | WGS | genomic | 101 |
| *Artemisia annua* | ERR11535563 | illumina NovaSeq 6000 | paired | WGS | genomic | 150 |
| *Bidens hawaiensis* | SRR14191093 | pacBIO SMRT | single | WGS | genomic | - |
| *Carthamus tinctorius* | SRR2154065 | illumina HiSeq 1500 | paired | WGS | genomic | 101 |
| *Chrysanthemum indicum* | CRR389876 (genome warehouse) | illumina NovaSeq 6000 | paired | WGS | genomic | 150 |
| *Conyza canadensis* | NA | NA | NA | NA | NA | NA |
| *Glebionis coronaria* | SRR20302831 | illumina NovaSeq 6000 | paired | WGS | genomic | 150 |
| *Helianthus annuus* | SRR2919251 | illumina HiSeq 2000 | paired | WGS | genomic | 100 |
| *Helichrysum umbraculigerum* | ERR10735438 | pacBIO SMRT | single | WGS | genomic | - |
| *Mikania micrantha* | SRR8835137 | illumina HiSeq X Ten | paired | WGS | genomic | 150 |
| *Pluchea indica* | SRR18449574 | illumina NovaSeq 6000 | paired | RNA-Seq | transcriptomic | 150 |
| *Scalesia atractyloides* | ERR9715097 | pacBIO SMRT | single | WGS | genomic | - |
| *Smallanthus sonchifolius* | SRR18215734 | illumina NovaSeq 6000 | paired | WGS | genomic | 150 |
| *Stevia rebaudiana* | SRR6792730 | illumina HiSeq X Ten | paired | WGS | genomic | 151 |
| *Tagetes patula* | SRR19579335 | illumina HiSeq X Ten | paired | WGS | genomic | 150 |
| *Tanacetum cinerariifolium* | SRR17714824 | illumina NovaSeq 6000 | paired | WGS | genomic | 151 |

**data for outgroups**

| species | read accession | platform | library layout | library strategy | library source | read length |
|---|---|---|---|---|---|---|
| *Beta vulgaris* | SRR868931 | Roche 454 GS FLX Titanium | paired | WGA | genomic | 100 |
| *Fragaria x ananassa* | SRR8358385 | illumina HiSeq 4000 | paired | WXS | genomic | 150 |
| *Lotus japonicus* | DRR014730 | illumina HiSeq 2000 | paired | WGS | genomic | 100 |
| *Hordeum vulgare* | SRR1804518 | illumina HiSeq 2000 | paired | WGS | genomic | 90 |

### Suppl. table 1E: Data sources and metadata for linkage vizualisation

**data for Asteraceae species**

| species | genome_size* [Gbp] | genome_size_ref | DOI | read_accession | cov_used | seq_technique |
|---|---|---|---|---|---|---|
| *Arctium lappa* | 1.79 | Song et al., 2023 | 0.1186/s12870-023-04092-3 | ERR5554584 | 0.5x | illumina |
| *Artemisia annua* | 1.74 | Shen et al., 2018 | 10.1016/j.jplph.2018.11.007 | (Upload needed) | 1x | illumina |
| *Bidens hawaiensis* | 7.56 | Bellinger et al, 2022 | 10.1093/jhered/esab077 | SRR14191093 | 1x | PacBio HiFi |
| *Carthamus tinctorius* | 1.32 | Nasab et al., 2023 | 10.1038/s41598-023-33347-0 | SRR2154065 | 1x | illumina |
| *Chrysanthemum indicum* | 3.02 | Li et al., 2012 | 10.1111/j.1759-6831.2012.00241.x | CRR389876 | 1x | ONT |
| *Glebionis coronaria* | 6.80 | Wang et al., 2022 | 10.1093/dnares/dsac036 | SRR20302831 | 1x | illumina |
| *Helianthus annuus* | 3.60 | Staton et al., 2012 | 10.1111/j.1365-313X.2012.05072.x | SRR2919251 | 0.3x | illumina |
| *Helichrysum umbraculigerum* | 1.30 | CATIUR000000000.1 | NA | ERR10735438 | 1x | illumina |
| *Mikania micrantha* | 1.87 | Liu et al., 2020 | 10.1186/s12864-019-6361-7 | SRR8835137 | 1x | illumina |
| *Scalesia atractyloides* | 3.2 | Cerca, 2022 | 10.5061/dryad.8gtht76rh | ERR9715097 | 1x | PacBio HiFi |
| *Smallanthus sonchifolius* | 2.72 | Fan et al., 2022 | 10.1111/1755-0998.13675 | SRR18215734 | 1x | illumina |
| *Stevia rebaudiana* | 0.40 | O'Neill and Pirro, 2020 | 10.12688/f1000research.24396.1 | SRR6792730 | 1x | illumina |
| *Tagetes patula* | 0.43 | JAOYNQ000000000.1 | NA | SRR19579335 | 1x | illumina |
| *Tanacetum cinerariifolium* | 7.10 | Yamashiro et al 2019 | 10.1038/s41598-019-54815-6 | SRR17714824 | 1x | illumina |

**data for outgroups**

| | | | | | | |
|---|---|---|---|---|---|---|
| *Beta vulgaris* | 0.75 | Dohm et al., 2014 | 10.1038/nature12817 | SRR952964 | 1x | illumina |
| *Fragaria x ananassa* | 0.70 | Isobe et al., 2018 | 0.1007/978-3-319-76020-9_10 | SRR16002690 | 1x | illumina |
| *Lotus japonicus* | 0.50 | Mun et al., 2016 | 10.1038/srep39447 | DRR014730 | 1x | illumina |
| *Hordeum vulgare* | 5.10 | Hisano et al., 2016 | 10.1186/s12864-016-3159-3 | SRR21763618 | 1x | illumina |

**** measured or assembled genome size as used for coverage calculation***
