## Supplemental Table 2: Cassandra and 5S metadata for "Evolving together: Cassandra retrotransposons gradually mirror promoter mutations of the 5S rRNA genes"

**Suppl. table 2A: Properties of all Cassandra retrotransposons within the dataset.** Lengths are goven in bp and identical sites in LTR regions in %. Sequence motifs for primer binding site (PBS) and polypurin tract (PPT) are also provided.

| order | plant family | species | length [bp] | 5' LTR [bp] | 3' LTR [bp] | internal region [bp] | LTR identical sites [%] | PBS | PPT |
|---|---|---|---|---|---|---|---|---|---|
| Cyatheales | Cyatheaceae | *Sphaeropteris cooperi* | 606 | 208 | 205 | 193 | 100.0 | TGGTATCAGAGC | GGGGCGGTTGT |
| Polypodiales | Didymochlaenaceae | *Didymochlaena trunculata* | 606 | 208 | 208 | 190 | 100.0 | TGGTATCAGAGC | AAGGGGGCGAT |
| | Nephrolepidaceae | *Nephrolepis exaltata* | 565 | 187 | 187 | 191 | 100.0 | TGGTATCAAAGC | AAGGGGGCGAT |
| Piperales | Aristolochiaceae | *Saruma henryi* | 563 | 234 | 234 | 95 | 100.0 | TGGTATCAGAGC | GAGGGGGTGAT |
| Poales | Poaceae | *Ambylopyrum muticum* | 724 | 264 | 264 | 196 | 100.0 | TGGTATGAGAGC | GTGGGGGTGTA |
| | Poaceae | *Avena sativa* | 773 | 287 | 286 | 200 | 99.7 | TGGTATCAGAGC | GGGTGGGCGTA |
| | Poaceae | *Brachypodium distachyon* | 668 | 251 | 251 | 166 | 100.0 | TGGTATCAGAGC | GGAGGGGTGTG |
| | Poaceae | *Bromus sterilis* | 689 | 266 | 266 | 157 | 100.0 | TGGTATCAGAGC | GTGGGGGTGTA |
| | Poaceae | *Colpodium drakensbergense* | 731 | 283 | 283 | 165 | 100.0 | TGGTATCAGAGC | GGGGGGAAGGA |
| | Poaceae | *Colpodium versicolor* | 729 | 282 | 282 | 165 | 100.0 | TGGTATCAGAGC | GGGGGGAAGGA |
| | Poaceae | *Deschampsia antarctica* | 766 | 283 | 283 | 200 | 100.0 | TGGTATCAGAGC | GGGGGGGTGTG |
| | Poaceae | *Eremopyrum distans* | 724 | 264 | 264 | 196 | 100.0 | TGGTATCAGAGC | GTGGGGGTGTA |
| | Poaceae | *Henrardia persica* | 726 | 264 | 264 | 198 | 100.0 | TGGTATCAGAGC | GTGGGGGTGTA |
| | Poaceae | *Hordeum brachyantherum* | 734 | 268 | 268 | 198 | 100.0 | TGGTATCAGAGC | GTGGGGGTGTA |
| | Poaceae | *Hordeum marinum* | 725 | 264 | 264 | 197 | 100.0 | TGGTATCAGAGC | GTGGTGGTGTA |
| | Poaceae | *Hordeum vulgare* | 725 | 264 | 264 | 197 | 100.0 | TGGTATCAGAGC | GTGGTGGTGTA |
| | Poaceae | *Oryza brachyantha* | 752 | 276 | 275 | 201 | 92.1 | TGGTATCAGAGC | GGGTGGGTGTA |
| | Poaceae | *Oryza glaberrima* | 768 | 281 | 281 | 206 | 100.0 | TGGTATCAGAGC | GGGGGTGAGGG |
| | Poaceae | *Oryza minuta* | 765 | 281 | 281 | 203 | 100.0 | TGGTATCAGAGC | GGGGGTGAGGA |
| | Poaceae | *Oryza sativa indica* | 766 | 280 | 280 | 206 | 99.6 | TGGTATCAGAGC | GGGGGTGAGGG |
| | Poaceae | *Oryza sativa japonica* | 768 | 281 | 281 | 206 | 99.3 | TGGTATCAGAGC | GGGGGTGAGGG |
| | Poaceae | *Panicum virgatum* | 869 | 280 | 280 | 309 | 100.0 | TGGTATCAGAGC | GGGGGTGAGCA |
| | Poaceae | *Peridictyon sanctum* | 732 | 268 | 268 | 196 | 100.0 | TGGTATCAGAGC | GTGGGGGTGTA |
| | Poaceae | *Phleum pratense* | 734 | 284 | 284 | 166 | 98.6 | TGGTATCAGAGC | GGGGGGAAGGA |
| | Poaceae | *Psathyrostachys fragilis* | 731 | 268 | 268 | 195 | 100.0 | TGGTATCAGAGC | GTGGGGGTGTA |
| | Poaceae | *Saccharum hybrid* | 747 | 273 | 273 | 201 | 100.0 | TGGTATCAGAGC | GGGGGTGAGGA |
| | Poaceae | *Secale cereale* | 724 | 264 | 264 | 196 | 100.0 | TGGTATCAGAGC | GTGGGGGTGTA |
| | Poaceae | *Setaria italica* | 767 | 281 | 282 | 204 | 96.8 | TGGTATCAGAGT | GGGGGTGAGTA |
| | Poaceae | *Spartina alterniflora* | 813 | 277 | 277 | 259 | 100.0 | TGGTATCAGAGC | GGGGGGTGGAT |

| | | | | | | | | | |
|---|---|---|---|---|---|---|---|---|---|
| | Poaceae | *Sorghum bicolor* | 742 | 288 | 288 | 166 | 89.9 | TGGTATCAGAGC | GGGGGTGAGGA |
| | Poaceae | *Triticum aestivum* | 724 | 264 | 264 | 196 | 100.0 | TGGTATCAGAGC | GTGGGGGTGTA |
| | Poaceae | *Zea mays* | 758 | 280 | 280 | 198 | 100.0 | TGGTATTAGAGC | AAAGGAGGGAT |
| | Poaceae | *Zingeria biebersteiniana sp. trichopoda* | 731 | 283 | 283 | 165 | 100.0 | TGGTATCAGAGC | GGGGGGAAGGA |
| | Poaceae | *Zingeria biebersteiniana* | 731 | 283 | 283 | 165 | 100.0 | TGGTATCAGAGC | GGGGGGAAGGA |
| | Poaceae | *Zingeria kochii* | 729 | 282 | 282 | 165 | 100.0 | TGGTATCAGAGC | GGGGGGAAGGA |
| | Poaceae | *Zingeria pisidica* | 729 | 282 | 282 | 165 | 100.0 | TGGTATCAGAGC | GGGGGGAAGGA |
| Malpighiales | Clusiaceae | *Garcinia mangostana* | 949 | 439 | 439 | 71 | 100.0 | TGGTATCAGAGC | GCTGGTGGGCA |
| | Euphorbiaceae | *Jatropha curcas* | 830 | 377 | 379 | 74 | 89.2 | TGGTATCAGAGC | GCTAGTGGGCC |
| | Linaceae | *Linum usitatissimum* | 632 | 276 | 271 | 85 | 100.0 | TGGTATCCGAGC | GGGGGGTGTAA |
| Fabales | Fabaceae | *Cajanus cajan* | 922 | 434 | 415 | 73 | 79.5 | TGGTATCAGAGC | GTTGGTGGGCT |
| | Fabaceae | *Glycine max* | 968 | 444 | 452 | 72 | 79.6 | TGGTATCATAAC | GCTGGTGGGCA |
| | Fabaceae | *Lens culinaris* | 912 | 416 | 414 | 82 | 99.8 | TGGTATCAGAGC | TTGGTGGGCCA |
| | Fabaceae | *Lotus japonicus* | 856 | 393 | 393 | 70 | 98.7 | TGGTATCAGAGC | GCAGGTGGGCC |
| | Fabaceae | *Medicago truncatula* | 858 | 382 | 406 | 70 | 87.9 | TGGTATAAGAGC | GCTGGTGGACA |
| | Fabaceae | *Pisum sativum* | 913 | 421 | 421 | 71 | 100.0 | TGGTATCAGAGC | GCTGGTGGGCA |
| Rosales | Cannabaceae | *Cannabis sativa* | 908 | 431 | 406 | 71 | 83.4 | TGGTATTACAGC | AGGGAGTTGAT |
| | Rosaceae | *Chaenomeles japonica* | 665 | 297 | 297 | 71 | 100.0 | TGGTATCAGAGC | AGGGGGTGGAT |
| | Rosaceae | *Fragaria x ananassa* | 609 | 267 | 267 | 75 | 100.0 | TGGTATCAGAGC | GGGGGGTGGAT |
| | Rosaceae | *Malus domestica* | 643 | 286 | 286 | 71 | 100.0 | TGGTATCAGAGC | GGGGGGTGGAT |
| | Rosaceae | *Prunus domestica* | 615 | 270 | 270 | 75 | 100.0 | TGGTATCAGAGC | GGGGGGTGGAT |
| | Rosaceae | *Rosa hybrid* | 669 | 299 | 299 | 71 | 100.0 | TGGTATCAGAGC | GGGGGGTGGAT |
| | Rosaceae | *Rosa rugosa* | 670 | 299 | 299 | 72 | 100.0 | TGGTATCAGAGC | GGGGGGTGGAT |
| | Rosaceae | *Rubus idaeus* | 669 | 300 | 299 | 70 | 100.0 | TGGTATCAGAGC | GGGGGGTGGAT |
| Brassicales | Brassicaceae | *Arabidopsis lyrata* | 831 | 361 | 359 | 111 | 96.7 | TGGTATCAGAGC | GGGGGGTGAAT |
| | Brassicaceae | *Arabidopsis thaliana* | 824 | 356 | 356 | 112 | 100.0 | TGGTATCAGAGC | TGTGGGTGAAT |
| | Brassicaceae | *Brassica oleracea* | 805 | 350 | 350 | 105 | 100.0 | TGGTATCAGAGC | AGGGGGTGAAT |
| | Brassicaceae | *Brassica rapa* | 803 | 349 | 349 | 105 | 100.0 | TGGTATCAGAGC | AGGGGGTGAAT |
| | Brassicaceae | *Thellungiella parvula* | 778 | 344 | 344 | 90 | 81.8 | TGGT-TCGGAGC | TATGGGTGAAT |
| | Brassicaceae | *Thellungiella salsuginea* | 778 | 344 | 344 | 90 | 81.8 | TGGT-TCGGAGC | TATGGGTGAAT |
| Caryophyllales | Aioaceae | *Mesembryanthemum crystallinum* | 665 | 300 | 300 | 65 | 100.0 | TGGTATCAGAGC | CTGGTCAGCCC |
| | Amaranthaceae | *Amaranthus palmeri* | 659 | 271 | 271 | 117 | 100.0 | TGGTATCAGAGC | GTGGGGGTGAA |
| | Amaranthaceae | *Beta vulgaris* | 761 | 283 | 283 | 195 | 95.4 | TGGTATTAGAGC | GTGGGGGTGTA |

| | | | | | | | | | |
|---|---|---|---|---|---|---|---|---|---|
| | Amaranthaceae | *Chenopodium quinoa* | 791 | 290 | 292 | 209 | 88.4 | TGATATCAGAGC | GTGGGGGTGTA |
| | Caryophyllaceae | *Colobanhus quitensis* | 801 | 302 | 302 | 197 | 100.0 | TGATATCAGAGC | GTGGGGGTGAT |
| | Caryophyllaceae | *Silene latifolia* | 806 | 315 | 315 | 176 | 96.8 | TGGTATCAAAGC | TGGGGGGGAAT |
| Ericales | Ericaceae | *Vaccinium corymbosum* | 650 | 235 | 235 | 180 | 100.0 | TGGTATCAGAGC | GGTGGGGAGAA |
| Asterales | Asteraceae | *Arctium lappa* | 647 | 283 | 283 | 81 | 100.0 | TGGTATCAGAGC | GGGGGGGTGTT |
| | Asteraceae | *Artemisia annua* | 663 | 299 | 282 | 82 | 92.0 | TGGTATCAGAGC | AGGGGGGTGAT |
| | Asteraceae | *Bidens hawaiensis* | 706 | 270 | 268 | 168 | 93.0 | TGGTATCAGAGC | GAGGGGGTGTT |
| | Asteraceae | *Carthamus tinctorius* | 616 | 269 | 266 | 81 | 96.7 | TGGTATCAGAGC | GGGGGGGTGTT |
| | Asteraceae | *Chrysanthemum indicum* | 623 | 269 | 267 | 87 | 86.6 | TGGTATCAGAGC | GGGGGGTGAAT |
| | Asteraceae | *Conyza canadensis* | 594 | 255 | 250 | 89 | 82.1 | TGGTATCCAAGC | TTGAGAGGGTG |
| | Asteraceae | *Glebionis coronaria* | 630 | 266 | 266 | 98 | 95.5 | TGGTATCAGAGC | GAGGGGGTGAT |
| | Asteraceae | *Helianthus annus* | 738 | 328 | 328 | 82 | 96.0 | TTGTATCAGAGC | AGGGGGGTGAT |
| | Asteraceae | *Helichrysum umbraculigerum* | 623 | 263 | 263 | 97 | 98.5 | TGGTATCAAAGC | GGGGGGTGAGT |
| | Asteraceae | *Mikana micrantha* | 625 | 270 | 270 | 85 | 97.0 | TGGTATCAGAGC | GGGGGGGTGTA |
| | Asteraceae | *Pluchea indica* | 601 | 264 | 256 | 81 | 74.3 | TGGTATCAGAGC | ACGGGGGTGTA |
| | Asteraceae | *Scalesia atractyloides* | 741 | 328 | 331 | 82 | 92.1 | TGGTATCAGAGC | AGGGGGGTGAT |
| | Asteraceae | *Smallanthus sonchifolius* | 641 | 277 | 277 | 87 | 93.9 | TGGTATCAGAGC | GGGGGGGTATT |
| | Asteraceae | *Stevia rebaudiana* | 732 | 323 | 328 | 81 | 89.1 | TGGTATCAGAGC | AAGGGGGGTGT |
| | Asteraceae | *Tanacetum cinerariifolium* | 613 | 266 | 266 | 81 | 88.0 | TGGTATCAGGGC | AAGGGGGTGAT |

**Suppl. table 2B: Position of 5S similarity regions in Cassandra LTR sequences.** Comparison was only done for species with a corresponding 5S rRNA gene. A-Box and C-Box motifs within the Cassandra are provided and nucleotides defininf the two main variants are highlighted in bold. Start and stop of the similarity regions are defined by the corresponding nucleotid position. The (rounded) relative values for 5S rDNA similariy region position are given in %.

| species | LTR length [bp] | A-Box | C-Box | 5S similiarity [nt] | similiarity start | similiarity stop | range position [%]* |
|---|---|---|---|---|---|---|---|
| *Amblyopyrum muticum* | 264 | AGTTAAGCGTGC | AG**GA**TGGGTG | 80 | 71 | 151 | 27 - 57 |
| *Avena sativa* | 287 | GGTTAAGCGTGC | AG**GA**TGGGTG | 72 | 80 | 151 | 29 - 53 |
| *Brachipodium distachyon* | 251 | AGTTAAGCATGC | AG**GA**TGGGTG | 96 | 62 | 157 | 25 - 63 |
| *Eremopyrum distans* | 264 | AGTTAAGCGTGC | AG**GA**TGGGTG | 96 | 66 | 161 | 25 - 61 |
| *Henrardia persica* | 264 | AGTTAAGCGTGC | AG**GA**TGGGTG | 81 | 63 | 143 | 24 - 54 |
| *Hordeum brachyantherum* | 268 | AGTTAAGCGTGC | AG**GA**TGGGTG | 73 | 73 | 145 | 27 - 54 |
| *Hordeum marinum* | 264 | GGTTAAGCGTGC | AG**GA**TGGGTG | 78 | 67 | 144 | 25 - 55 |
| *Hordeum vulgare* | 264 | GGTTAAGCGTGC | AG**GA**TGGGTG | 66 | 80 | 145 | 30 - 55 |
| *Oryza brachyantha* | 276 | GGTTAAGCGTGT | GG**GA**TGGGTG | 87 | 70 | 155 | 25 - 56 |
| *Oryza sativa* | 281 | GGTTAAGCGTGC | AG**GA**TGGGTG | 75 | 80 | 154 | 28 - 55 |
| *Panicum virgatum* | 280 | GGTTAAGCGTGC | GG**AT**GGGTGA | 86 | 70 | 155 | 25 -55 |
| *Setaria italica* | 281 | GGTTAAGCGTGC | GG**GA**TTGGTG | 86 | 70 | 155 | 25 - 55 |
| *Secale cereale* | 264 | AGTTAAGCGTGC | AG**GA**TGGGTG | 79 | 66 | 144 | 25 - 55 |
| *Triticum aestivum* | 264 | AGTTAAGCGTTC | AG**GA**TGGGTG | 72 | 73 | 144 | 28 - 55 |
| *Zea mays* | 280 | AGTTAAGCGTGC | GG**GA**TGGGTG | 74 | 88 | 161 | 31 - 58 |
| *Linum usitatissimum* | 276 | TGTTAATCGCGC | TA**AA**TGGGTG | 97 | 108 | 204 | 39 - 74 |
| *Jatropha curcas* | 379 | GGTTAAGCGTGC | AG**GA**TGGGTG | 78 | 188 | 265 | 50 - 70 |
| *Glycine max* | 444 | AGTTAAGCGTGC | GA**GA**TGGGTG | 87 | 132 | 218 | 30 - 49 |
| *Lotus japonicus* | 395 | AGTTAAGTGTGC | AG**GA**TGGGTG | 73 | 238 | 310 | 60 - 78 |
| *Medicago truncatula* | 382 | AGTTAAGCGTGC | GG**GA**TGGGTG | 75 | 217 | 291 | 57 - 76 |
| *Pisum sativum* | 421 | AGTTAAGCGTGC | GG**GA**TGGGTG | 69 | 256 | 329 | 61 - 78 |
| *Cannabis sativa* | 431 | GGTTAAGCGTGC | TG**GG**TGGATG | 77 | 194 | 270 | 45 - 63 |
| *Fragaria x ananassa* | 267 | GGTTAAGCATGT | AG**GA**TGGGTG | 84 | 96 | 179 | 36 - 67 |
| *Malus domesticus* | 286 | AGTTAAGCGAGA | AT**GA**TGGGTG | 104 | 78 | 181 | 27 - 63 |
| *Arabidopsis lyrata* | 359 | AGTTAAGCGTGC | AG**GA**TGGGTG | 86 | 106 | 193 | 30 - 54 |
| *Arabidopsis thaliana* | 356 | AGTTAAGCGTGC | AG**GA**TGGGTG | 80 | 104 | 182 | 29 - 51 |
| *Brassica rapa* | 349 | AGTTAAGTGTGC | AG**GA**TGGGTG | 72 | 92 | 163 | 26 - 47 |
| *Beta vulgaris* | 283 | AGTTAAGTGTGC | GG**GA**TGGGTG | 70 | 90 | 159 | 32 - 56 |

| | | | | | | | |
|---|---|---|---|---|---|---|---|
| *Silene latifolia* | 315 | AGTTAAGCGTGC | AT**GA**TGGGTG | 79 | 113 | 191 | 36 - 61 |
| *Arctium lappa* | 283 | GGTTAAGCGTGC | AG**GA**TGGGTG | 68 | 122 | 189 | 43 - 67 |
| *Artemisia annua* | 282 | AGTTAAGCGTGC | GG**CC**TGGGTG | 66 | 118 | 184 | 42 - 65 |
| *Bidens hawaiensis* | 270 | AGTTAAGCGTGC | AG**AT**GGTTGA | 68 | 131 | 198 | 49 - 73 |
| *Carthamus tinctorius* | 269 | AGTTAAGCGTGC | AG**GA**TGGGTG | 68 | 124 | 191 | 46 - 71 |
| *Chrysanthemum indicum* | 269 | AGTTAAGCATGC | AG**CC**TGGGTG | 67 | 132 | 198 | 49 - 74 |
| *Conyza canadensis* | 255 | AGTTAAGCGTGT | CT**TA**TGGGTG | 66 | 121 | 187 | 47 - 73 |
| *Glebionis coronaria* | 266 | AGTTAAGCGTGC | GC**TT**GGGTGA | 74 | 126 | 192 | 47 - 72 |
| *Helianthus annuus* | 328 | AATTAAGTGTGC | AG**GA**TGGGTG | 90 | 154 | 245 | 47 - 75 |
| *Helichrysum umbaculigerum* | 263 | AGTTAAGCGTGC | AG**GA**TGGGTG | 65 | 127 | 191 | 42 - 73 |
| *Mikana micrantha* | 270 | AGTTAAGCGTGC | GG**GA**TGGGTG | 91 | 110 | 200 | 41 - 74 |
| *Pluchea indica* | 264 | AGTTAAGCGTGC | TG**GA**TATGTG | 77 | 111 | 187 | 42 - 71 |
| *Scalesia atractyloides* | 328 | AGTTAAGCGTGC | GG**AT**GGGTGA | 67 | 175 | 249 | 53 - 76 |
| *Smallanthus sonchifolius* | 277 | AGTTAAGCGTGC | GG**GA**TAGGTG | 67 | 139 | 196 | 50 - 71 |
| *Stevia rebaudiana* | 323 | AGTTAAGCGTGA | AG**GA**TGGGTG | 84 | 159 | 242 | 49 - 75 |
| *Tanacetum cinerariifolium* | 266 | AGTTAAGCGTGC | GG**CC**TGGGTG | 71 | 117 | 187 | 44 - 70 |

****  calclated ratios are rounded***
